# PlantMetWiki: a FAIR knowledge graph for plant metabolic pathway cross-species representation and integration

**DOI:** 10.64898/2026.07.22.733699

**Authors:** Elena Del Pup, Max Muller, Marvin Martens, Egon L. Willighagen, Marnix H. Medema, Denise Slenter, Justin J.J. van der Hooft

## Abstract

Plants produce a vast diversity of specialized metabolites with extensive potential ecological, agro-industrial, and pharmaceutical applications. Discovery of novel plant natural products relies on combining multi-omics evidence with biochemical transformations. However, pathway-level annotations are fragmented across individual species, databases, and publications, limiting comparative cross-species pathway analyses and systematic generation of hypotheses. To support integration and reuse of plant metabolic knowledge, we developed PlantMetWiki, a FAIR Linked Open Data semantically enriched knowledge graph built on infrastructure adapted from WikiPathways. Our approach extends on the highly curated pathway information from Plant Metabolic Network with crosslinks to biosynthetic gene clusters resources (MIBiG and plantiSMASH), metabolite annotations, and cross-species modelling. This way, our resource captures pathway genomic context, increases metabolomics data interoperability via federated queries, and supports cross-species analysis to identify annotation gaps. PlantMetWiki represents 1,162 plant metabolic pathways as Resource Description Framework (RDF) graphs, preserving pathway structure, literature provenance, annotations, and taxonomic information from PlantCyc 17.0. PlantMetWiki is distributed through a public SPARQL endpoint with open-source reproducible data transformation and validation workflows. By modelling pathways as a multispecies graph, PlantMetWiki enables comparative analyses across taxa, integration with external chemical knowledge resources through federated queries, and identification of metabolic, genomic, and chemical annotation gaps. As a result, PlantMetWiki provides a foundation for FAIR reuse and integration of plant pathway knowledge.

## Introduction

Plants encompass hundreds of thousands of species that have evolved diverse metabolic repertoires to adapt to distinct ecological niches ^1^. As a result, different plant lineages produce a vast number of unique combinations of endogenous specialized metabolites, ranging from alkaloids and terpenoids to phenylpropanoids and glycosides, many of which have important ecological, agricultural, industrial, and pharmaceutical applications ^2–4^. As evolution of plant chemical diversity largely relies on shared metabolic networks and common branchpoints ^5^, mapping biosynthetic pathways across species remains a central challenge in plant biology and natural product research.

While the increasing number of experimentally characterized plant metabolic pathways provides valuable insights into enzymes, metabolites, and regulatory mechanisms underlying specialized metabolism, biosynthetic knowledge remains unevenly distributed across species, pathways, and specialized domain-specific resources. Model organisms and economically important crops are extensively annotated; however, many species contain only partial pathway information, and evidence for individual pathway steps is often scattered across multiple taxa. Existing databases capture complementary yet disconnected aspects of metabolism, including enzymatic reactions, metabolite structures, or genomic context, but are rarely integrated. While these resources are individually valuable, they are rarely represented within a common framework to allow pathway, genomic, taxonomic, and chemical information to be explored simultaneously. For instance, major natural product chemical structure databases such as LOTUS ^6^ and COCONUT ^7^ contain structures and associated metadata but lack explicit links to metabolic pathways, thereby hindering cross-species analyses supporting enrichment of multi-omics datasets with metabolite information or annotation of cross-species metabolic networks ^8,9^. Integrating this fragmented knowledge is essential for comparative pathway analysis, biosynthetic hypothesis generation, and the discovery of novel plant natural products. Comparative analyses across species are also essential to translate knowledge from model plant species to less-studied taxa and crops ^10^, as well as to understand the evolutionary processes that shape specialized metabolism. Pathways may diversify through lineage-specific innovations, recruit alternative enzymes to catalyze equivalent reactions, or arise independently through convergent evolutionary strategies that lead to the production of similar metabolites in distantly related species ^11^. The relationships between species are captured by taxonomic information, where NCBI Taxonomy ^12^ acts as a reference ontology. Despite the importance of these comparative perspectives, most pathway resources and their semantic representation remain centered on species-specific representations, limiting the ability to systematically query shared pathway components and integrate evidence across biological and omics layers. Cross-species pathway models are a promising framework to connect these different biological entities together in a harmonized overview with many options for data visualization, analysis, and interpretation.

The Plant Metabolic Network (PMN) ^13^ provides the extensively curated pan-plant reference database PlantCyc, which includes machine-readable annotations of enzymes and metabolites across species-specific biochemical conversions across hundreds of plant taxa. PMN adopts the BioPAX ^14^ standard to facilitate pathway data exchange and interoperability and the Pathway Tools ecosystem ^15^. This standardized format, however, presents limitations for integrative analyses, since the flat tabular representations simplify access at the cost of losing the rich relational structure of metabolic networks. Consequently, complex relationships such as hierarchical pathway organization, many-to-many associations between reactions and metabolites, and detailed provenance information are difficult to represent and query. Furthermore, this format cannot accommodate integration with other annotations, including biosynthetic gene clusters, predicted pathways, individual multi-omics datasets, and external knowledge graphs, restricting large-scale comparative and cross-domain analyses.

The Graphical Pathway Markup Language (GPML) ^16^ provides a pathway modeling standard that is more flexible in nature and can be extended with new annotations. This format can be transferred to the Resource Description Framework (RDF), providing an open, community-curated platform for pathway representation that adopts Semantic Web technologies using vocabularies enabling data sharing and reuse, persistent identifiers, and SPARQL for querying ^17^. This graph-based approach enables pathways to be represented as interconnected networks, preserving complex biological relationships while supporting integration with external resources through Linked Open Data (LOD) for machine-readable interlinked data. The infrastructure developed by WikiPathways ^18^ facilitates visualization, editing, and analysis of pathway networks, and supports integration with multi-omics tools and network analysis platforms such as Cytoscape ^19^, as well as federated queries across databases including Wikidata ^20^. In plants, the use of WikiPathways was piloted to analyze rice and Arabidopsis seed development networks and gene regulatory networks ^21^. However, the current GPML pathway model used in WikiPathways is constrained to single-species pathway representations, thereby limiting integration approaches that encompass multiple species and the genome-scale annotations available in PMN.

Emerging evidence of the physical clustering of plant biosynthetic genes in the chromosome could be used to annotate gaps in plant metabolic knowledge by integrating these genomic regions called biosynthetic gene clusters (BGCs) with pathway models. MIBiG ^22^ is a repository of experimentally validated BGCs which relies on a minimum information standard for their annotation, continuously powered by collaborative community annotations, and contains 43 validated plant BGCs in version 4.0. However, MIBiG lacks representation of partially clustered plant pathways, which are abundant in plants across taxa ^23^. Additionally, the MIBiG schema does not capture multispecies information for BGCs that occur in several species or that are similar to each other, despite many key studies for their discovery and characterization rely on comparative approaches ^24,25^. This limits follow-up comparative studies on the relationships between clustered and unclustered biosynthesis. The plantiSMASH ^26^ database captures more than 30,000 predicted plant BGCs across more than 400 species, detected by mining plant genomes for enzyme domains associated with known biosynthetic “rules”. The database maps plantiSMASH-detected BGCs to validated ones from MIBiG and to other similar putative BGCs across species, facilitating cross-species analyses. However, it does not provide links of plantiSMASH-detected BGCs with wider metabolic networks.

To overcome the above-mentioned gaps, here we present Plant Metabolic Pathways Wiki (PlantMetWiki), a LOD knowledge graph for representing and exploring plant metabolic pathways across species. PlantMetWiki combines curated pathway knowledge from PlantCyc with the semantic infrastructure of WikiPathways ^18^, creating a unified graph-based representation of plant metabolism that preserves pathway structure while enabling interoperability with external resources. The resource integrates experimentally characterized pathway annotations from PlantCyc 17.0, taxonomic information from NCBI, metabolite identifiers through mappings via BridgeDb ^27,28^, crosslinks to biosynthetic gene cluster information from MIBiG 4.0 ^22^ and predicted clusters from plantiSMASH 2.0 ^26^, and natural product chemistry through federated queries to Wikidata.

PlantMetWiki enables users to (1) compare pathways across species, leveraging shared reactions, enzymes, substrates, and products, to identify annotation gaps, (2) explore connections between curated pathways and their genomic context, including predictions from plantiSMASH genome mining, and (3) calculate completeness between complementary data resources in the natural products space with federated queries. In this work, we describe the generation of PlantMetWiki, its FAIR (Findable, Accessible, Interoperable, and Reusable) data infrastructure, and example applications demonstrating cross-species pathway comparison and integration with plant natural product knowledge. By representing pathways as a multispecies semantic graph with (gene, metabolite, and taxa) identifier mapping and RDF-based data models available for querying, PlantMetWiki connects pathway information to complementary biological, taxonomic, and chemical knowledge available across the LOD ecosystem and supports query-driven analyses. Additionally, our resource provides an expandable semantic model and a reproducible framework that can be transferred to other kingdoms of life using multispecies models for pathway knowledge gap discovery, such as microbes and fungi.

## Results

PlantMetWiki is accessible through an open-access SNORQL User Interface (UI) (https://plantmetwiki.bioinformatics.nl) and a dedicated SPARQL endpoint (https://plantmetwiki.bioinformatics.nl/sparql), including diverse customizable example queries. A dedicated tutorial includes relevant documentation of the UI, endpoint, and example queries, supporting integration with user-provided data, including multi-omics data, phenotypes, federated queries to Wikidata, and BGCs (from MIBiG and plantiSMASH). Results can be exported in standard formats such as CSV, JSON, and XML to be further analyzed using network tools such as Cytoscape and combined with multi-omics visualizations and integrative representations such as metabolic networks ^8^ or paired omics approaches ^29^.

Querying the semantically modelled pathway information in PlantMetWiki supports identification of curation gaps and prioritization of candidate annotations through alignment of cross-species pathway information and independent knowledge resources, crosslinks in the knowledge graph (to MIBiG, NCBI, plantiSMASH), and federated queries. PlantMetWiki supports the FAIR principles by adopting RDF-based semantic representations, including standard ontologies, machine-readable metadata, and public data access mechanisms, described in more detail in Table 1.

**Table 1:** Adaptation of the FAIR principles in the PlantMetWiki resource.

| FAIR principle | Implementation in PlantMetWiki |
| --- | --- |
| F1: (Meta) data are assigned globally unique and persistent identifiers | 1. Uniform Resource Identifiers (URIs) resource identifiers for all data elements,<br>2. Resolvable Uniform Resource Locators (URLs) for publications and metabolites for all species<br>3. Identifier mapping of genes and proteins with Resolvable Uniform |
|  | Resource Locators (URLs) in<br>*Arabidopsis thaliana* |
| F2. Data are described with rich metadata (defined by R1 below) | RDF model is supported with VoID header file, including license and provenance information. |
| F3. Metadata clearly and explicitly include the identifier of the data they describe | Identifier, version, and location of releases from PlantCyc, plantiSMASH, MiBIG, NCBI ontology, and BridgeDb metabolite mapping files are available in the VoID header file. |
| F4. (Meta)data are registered or indexed in a searchable resource | SPARQL endpoint |
| A1. (Meta)data are retrievable by their identifier using a standardized communications protocol<br>A1.1 The protocol is open, free, and universally implementable<br>A1.2 The protocol allows for an authentication and authorization procedure, where necessary | 1. Standardized access via a SPARQL endpoint<br>2. SPARQL documentation is open, free, and universally implementable, enhanced with a dedicated tutorial<br>3. Authentication and authorization can be implemented, however not necessary due to the nature of our data |
| A2. Metadata are accessible, even when the data are no longer available | 1. All data accessible via the SPARQL endpoint are archived on Zenodo<br>2. The software to create the data is available on GitHub |
| I1. (Meta)data use a formal, accessible, shared, and broadly applicable language for knowledge representation | 1. Framework is based on 10+years RDF model from WikiPathways<br>2. Void header file follows the Dublin Core Schema |
| I2. (Meta)data use vocabularies that follow FAIR principles | Included vocabularies are based on OWL representations available on BioPortal (WikiPathways) and OBO Foundry (NCBITaxon ontology) |
| I3. (Meta)data include qualified references to other (meta)data | Interactions between biological entities are described using the new GPML2021 |
|  | schema, including detailed biological interactions compatible with cross-references |
| R1. (Meta)data are richly described with a plurality of accurate and relevant attributes<br>R1.1. (Meta)data are released with a clear and accessible data usage license<br>R1.2. (Meta)data are associated with detailed provenance<br>R1.3. (Meta)data meet domain-relevant community standards | 1. License and provenance are recorded in the VoID header file for automated retrieval<br>2. License and provenance of data and software are included in the SNORQL UI banner, to support user awareness<br>3. GPML and BridgeDb are recognized resources in FAIRSharing, on which the PlantMetWiki tool is built |

### Building a cross-species semantic model and expandable RDF framework for plant specialized metabolism

The PlantMetWiki open-source web interface is the result of six co-designed and jointly operating repositories hosted in the GitHub Pathway-LOD community (see Code availability statement) that capture all steps from transforming data from PlantCyc in their native BioPAX standard to GPML2021 and RDF, metabolite mappings via BridgeDb, establishing BGC crosslinks, creating a metadata schema, versioning, automated, SNORQL UI hosting, SPARQL example querying, documentation, and tutorials (Figure 1). Data, metadata and software versioning and archiving on Zenodo are automated steps, with dedicated workflows available on GitHub (see Code availability statement). Pathway and reaction data from PlantCyc 17.0 are modelled into 1,162 multispecies pathway files and 1,316 individual enzymatic reaction files in the GPML2021 format, then converted to one RDF Turtle file using the WikiPathways gpml2rdf tool (v4.0.4), providing harmonized representations of all pathways and associated reactions. The pathways captured by both file types are represented as “wp:Pathway” objects and together constitute 2,478 pathway instances in the resulting core PlantMetWiki knowledge graph. Pathways contain different entity types (DataNodes), modelled following the WikiPathways vocabulary (http://vocabularies.wikipathways.org/wp#), representing 2,758 genes (wp:GeneProduct), 3,750 enzymes (wp:Protein), 70 protein complexes (wp:Complex), 4,577 metabolites (wp:Metabolite), and 54,800 Interactions (Fig. S1). Interactions encode directed biological events between DataNodes as “wp:target/source” edges (Figure 2), and are split into 15 subtypes, summarized in Fig. S2. Conversions (wp:Conversion) representing biochemical reactions involving metabolite substrates and products encode 36.4% (19,927) of the total Interactions, followed by Catalysis (wp:Catalysis), linking “wp:GeneProduct” or “wp:Protein” to the reaction these entities catalyze, with 22.6% (12,359), “wp:Transcription/Translation” (links genes, to the protein these encode) 15.1% (8,280), “wp:Inhibition/Stimulation” (negative or positive regulatory effects) 6.4% (3,486) and 1.6% (865), respectively, “wp:Binding”/ComplexBinding (physical interaction or complex formation) 0.1% (70). Some PlantCyc pathway edges, particularly transport and regulatory links, are translated to “wp:DirectedInteraction” (44,934, 82% total Interaction coverage) as a fallback type when no more specific class applies. The cumulative coverage of entities per pathway shows that a few large pathways dominate the resource, with genes and proteins cumulative curves across the entire database rising steeply (Fig. S3). Metabolites do not follow such a steep trend, since their nodes are often shared across pathways. The largest pathways in terms of genes, metabolites, and species represented from PlantCyc include super pathways of flavones and derivatives (PWY-6266), super pathway of anaerobic sucrose degradation (PWY-7345), and quercetin glycoside biosynthesis (PWY-5321) (Fig. S4-5).

**Figure 1:**
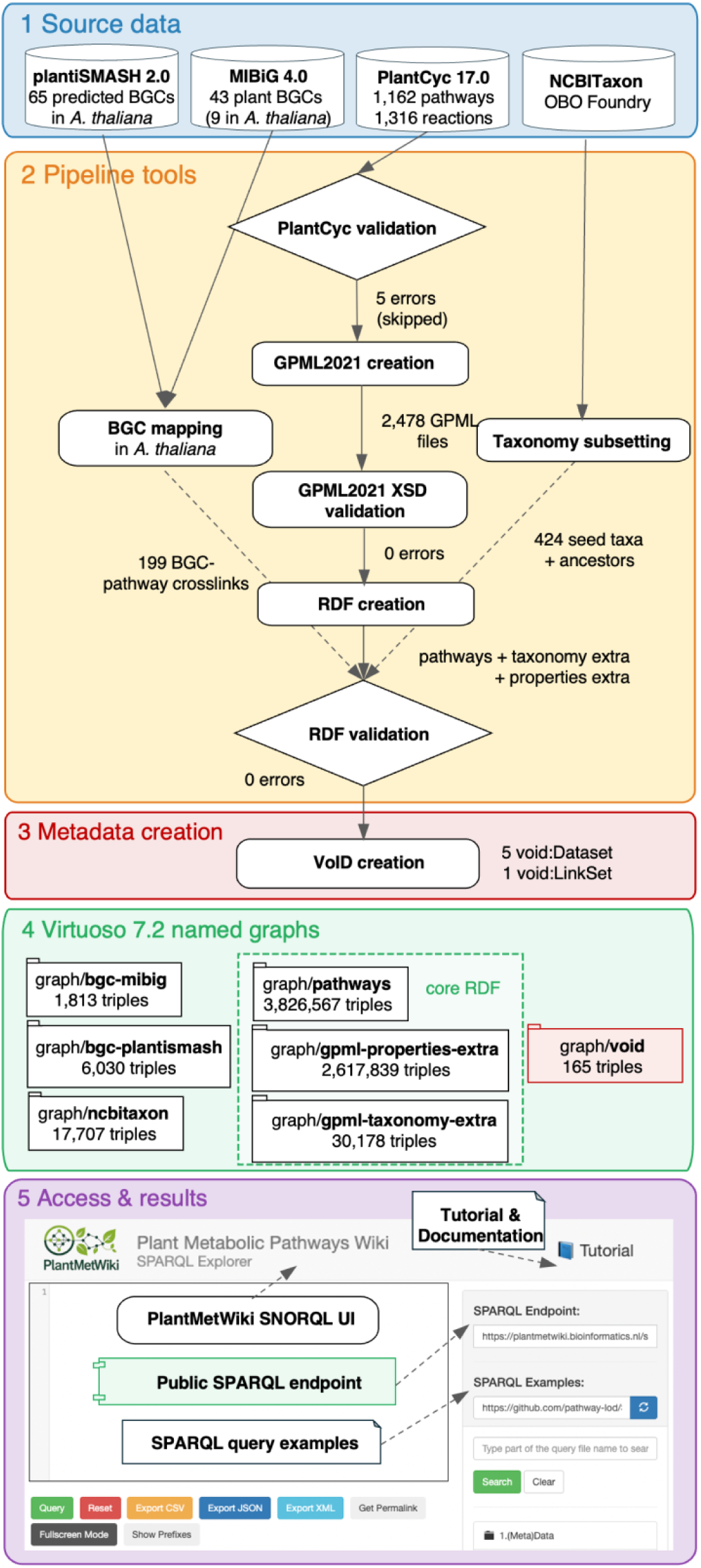
PlantMetWiki data transformation pipeline and architecture. All repositories for building and hosting PlantMetWiki are available on the pathway-LOD GitHub community (https://github.com/pathway-lod). The diagram shows the **five** stages of the PlantMetWiki knowledge-graph construction: **(1) Source data** (blue): MIBiG 4.0 and plantiSMASH 2.0 biosynthetic gene cluster (BGC) annotations, PlantCyc 17.0 biochemical databases (1,162 pathways, 1,316 reactions, 439 NCBI taxa), and the Open Bio-Ontologies (OBO) Foundry NCBITaxon ontology. NCBI taxonomy subset is generated with ROBOT v1.9.6 MIREOT extraction method. **(2) Pipeline tools** (yellow): PlantCyc input validation checks of proteins.dat and genes.dat for taxonomic annotation consistency before any files are generated (validate_plantcyc_input.py). The GPML conversion tool (build_pathways.py) produces 2,478 GPML2021 pathway and reaction files. A post-build XSD validator confirms schema compliance (test_gpml_files.py). The RDF conversion layer generates three named graphs (Java gpml2rdf-4.0.4.jar for main pathway graph; custom Python scripts). The BGC mapping “map-to-RDF" pipeline links BGC gene nodes to pathway enzymes. The RDF data is validated using a custom Python script that uses the RDFlib library. **(3) Metadata creation** (red): a VoID RDF Schema vocabulary is used to capture the metadata of all RDF datasets. This creation happens in three separate stages: core RDF conversion (create_void_from_metadata.py), BGC crosslinks mapping (create_void_bgc.py) and NCBI Taxonomy subsetting. **(4) Virtuoso triplestore** (green): Seven named graphs totaling ∼28.2 million triples. The pathway graphs hold the core WikiPathways RDF, entity-level species annotations, and PlantCyc-specific properties. The NCBITaxon graph (21.7M triples) provides taxon labels and hierarchy for all 439 NCBI taxa. **(5) Access and results** (purple): A public SPARQL endpoint (https://plantmetwiki.bioinformatics.nl/sparql) and SNORQL UI browser interface (https://plantmetwiki.bioinformatics.nl) allow direct querying of the Virtuoso named graphs. A tutorial and documentation (https://pathway-lod.github.io/SPARQLTutorials/) and SPARQL query examples are available (github.com/pathway-lod/SPARQLQueries). Jupyter notebooks support the creation of publication-ready figures, cross-species pathway inference analyses, and federated queries to external endpoints such as Wikidata. Solid arrows: automated steps; dashed arrows: manual steps.

**Figure 2:**
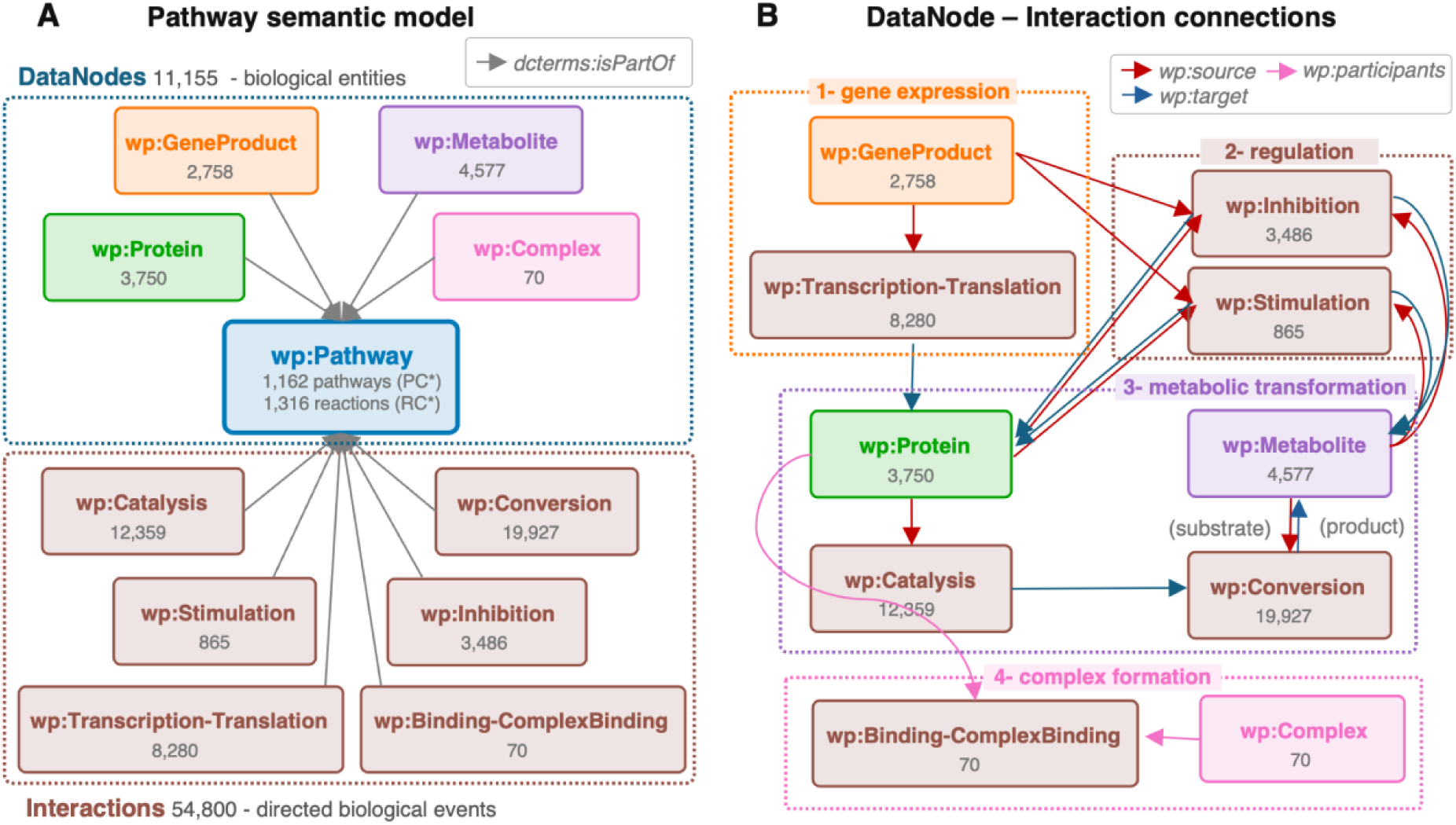
PlantMetWiki core graph model. **(A) Pathway semantic model.** Every pathway (wp:Pathway, blue centre; 1,162 PlantCyc pathways PC* and 1,316 reactions RC*) is associated with two categories of RDF resource via dcterms:isPartOf: DataNodes (biological entities; blue dashed box) and Interactions (directed biological events; brown dashed box). DataNode types: wp:GeneProduct (gene/transcript, n = 2,758 unique IRIs), wp:Protein (enzyme/protein, n = 3,750), wp:Metabolite (small molecule, n = 4,577), wp:Complex (n = 70). Interaction types with their total edge counts across all pathways: wp:Catalysis (12,359), wp:Conversion (19,927), wp:TranscriptionTranslation (8,280), wp:Inhibition (3,486), wp:Stimulation (865), wp:Binding/wp:ComplexBinding (70). Total interactions: 54,800. Total unique DataNodes: 11,155. **(B) DataNode-Interaction connection patterns,** Each wp:Interaction subtype connects two participants via wp:source (red arrows), wp:target (blue arrows), or wp:participants (pink arrows). The diagram shows all DataNode types that act as sources, targets, or participants for each Interaction type. Through the graph, four biological processes are represented: (1) Gene expression: wp:GeneProduct is the source of wp:TranscriptionTranslation which targets wp:Protein; (2) Regulation: wp:Inhibition and wp:Stimulation connect wp:GeneProduct, wp:Protein, or wp:Metabolites to the wp:Protein or wp:Metabolite target; (3) Metabolic transformation: wp:Catalysis takes a wp:Protein as source and targets a wp:Conversion node as target. wp:Conversion goes from one or more source wp:Metabolite (substrates) to one or more wp:Metabolite (products); (4) Complex formation: wp:Binding/wp:ComplexBinding links wp:Metabolite or wp:Complex nodes to a wp:Complex.

Taxonomic information from PlantCyc is modeled by mapping 455 PlantCyc-internal organism identifiers (ORG-IDs) to 439 unique NCBI Taxonomy identifiers. Sixteen ORG-codes representing different database instances of the same organism were collapsed to a single NCBI taxon ID. For instance, ORG-5993 (*Arabidopsis thaliana* ecotype Col-0 database) and TAX-3702 (*Arabidopsis thaliana*) both mapped to NCBI:3702 (Supplementary Table S1). Of the 439 NCBI taxa, 425 (96.8%) are connected to annotated DataNodes as Genes, Proteins, or Metabolites, with 98.7% of GeneProduct DataNodes and 98.8% of Protein DataNodes carrying taxon annotation (Fig. S1). By taxonomic rank, 414 are species-level binomial entries, 13 are interspecific hybrids or cultivar groups, 10 are genus-level annotations (where PlantCyc recorded the enzyme at genus rather than species resolution), and two remain unresolved (ARA and TAX-101007) (Supplementary Table S2). The remaining 12 taxa (2.7%) are present in PlantCyc exclusively as monomers of multi-subunit enzyme complexes (totaling only 0.1% coverage, 33 of 23,449 nodes). As the GPML2021 schema does not permit species annotations on <Group type="Complex"> elements, these taxa cannot be represented as DataNode annotations and are therefore absent from the species-level RDF output. For the two monomers that also appear as standalone DataNodes in at least one reaction or pathway context, this propagation successfully annotated two additional Protein DataNodes (from 10,583 to 10,585 total annotated Protein nodes). The remaining 10 monomers that are not annotated as direct reaction catalysts do not appear as standalone DataNodes in any GPML file and thereby do not receive a “wp:organism” annotation in the final RDF output. Of the 439 unique NCBI Taxonomy IDs represented in PlantCyc 17.0.0, 437 (99.5%) are annotated in the GPML output: 413 as Protein DataNodes, 306 as GeneProduct DataNodes, 24 as Metabolite DataNodes, and 12 exclusively via Complex Group elements. Two taxa remain absent: one whose proteins are not linked to any pathway or reaction in the GPML build (*Fragaria virginiana*, NCBITaxon:101015), and one with an unresolved NCBI Taxonomy identifier (TAX-101007).

The PlantCyc input validation pipeline identified five genes carrying annotations pointing to different NCBI taxa. For example, CYP93B2 (wp:GeneProduct G-9216) gene for flavone synthase II (wp:Protein MONOMER-12564) is annotated to both *Gerbera* hybrid cultivar (TAX-18101) and *Glycyrrhiza echinata* (TAX-46348), and PcFSI gene (wp:GeneProduct G-9647) for flavone synthase I (wp:Protein MONOMER-11863) is annotated to both *Petroselinum crispum* (TAX-4043) and *G. echinata* (TAX-46348). These cases point to a potential curation issue or inconsistency of the source database, which otherwise has unique gene identifiers for every taxon, and requires to be dealt with caution given that gene taxonomy annotation in the PlantMetWiki graph relies on direct annotation protein products. The validator reports this issue and skips taxonomy annotation for these five genes, to avoid erroneous links in the PlantMetWiki knowledge graph. The affected genes, their protein IDs, and source literature references are documented in Supplementary Table S3. Additionally, our custom validation pipeline captured two warning classes involving taxonomy of proteins and complexes (see Supplementary Table S3). First, 10 protein nodes were found to carry more than one taxonomic (SPECIES) value in PlantCyc: cases where the same enzyme has been biochemically characterized in multiple plant species (e.g., flavone synthase II characterized in both *G. echinata* and *Gerbera* hybrid cultivar). This issue is biologically valid in plants and is handled by our pipeline, where each listed taxon is written as a separate <AnnotationRef type="Taxonomy"> element on the protein DataNode and propagated to the encoding gene DataNode, ensuring all species annotations are preserved and modelled consistently in the downstream RDF graph. The second warning concerns one gene whose protein products carry different BioCyc ORG-codes that nevertheless resolve to the same NCBI taxon (e.g., ORG-5993 and TAX-3702 both map to NCBI:3702, *A. thaliana*), resulting in ORG-code mapping to their canonical NCBI taxon ID and annotation elementId (e.g., taxonomy_3702). A deduplication step when propagating taxonomic annotations from Proteins to Genes ensures the gene DataNode receives exactly one AnnotationRef per unique NCBI taxon, regardless of how many PlantCyc ORG-codes were listed.

Two informational log files were generated. First, for 28 genes that encode multiple protein products (isoforms), the species annotation of the gene DataNode is resolved by iterating over all isoform protein nodes and merging their taxon AnnotationRefs into one gene (deduplicated by elementRef), so every distinct taxon across all isoforms is preserved. Second, four protein entries list more than one “GENE” entry in proteins.dat (as is common for enzyme complexes or paralogous gene models). The species annotation is propagated to all listed genes, provided the corresponding gene records in genes.dat each carry the protein in their own “PRODUCT” field (see Supplementary Table S3) to avoid loss of information.

Species-level annotations in the final graph are encoded in a dedicated taxonomy graph (30,176 triples), which links individual gene and enzyme nodes to 424 distinct NCBITaxon identifiers covering 96% of gene product nodes (2,656/2,758) and 99% of enzyme nodes (3,712/3,750). When loading the full Open Bio-Ontologies (OBO) Foundry NCBITaxon release (v2026-05-13, 21,719,262 triples), six taxa from the 424 in total present in PlantCyc 17.0 were absent from the ontology, connected to 37 GeneProduct and Protein nodes from the modelled pathways that could not be resolved to a specific taxon label. This resulted in 418 taxa of the 424 original taxa being resolved to label via the OBO Foundry NCBITaxon ontology. From the six unmapped taxa, *Lycopersicon hirsutum* (NCBI:283673) has been deprecated and merged into *Solanum habrochaites* (NCBI:62890) in current NCBI Taxonomy releases, while the remaining five (*Dahlia variabilis, Foeniculum vulgare, Perilla citriodora, Schizonepeta tenuifolia,* and *Ailanthus altissima*) are valid NCBI taxa that are not included in this version of the OBO Foundry release. These 37 nodes represent 0.2% of the 18,762 species-annotated DataNodes in the knowledge graph and do not affect pathway structure or metabolite annotations (see Data availability). ROBOT ^30^ MIREOT extraction downsized the NCBITaxon ontology graph from 1.8GB of the 21.7 million triples of the entire graph to 1.4MB for a total of 17,707 triples capturing the whole lineage of the 424 represented taxa, a slimming mechanism required for hosting this data in the Virtuoso triplestore and avoiding common bottlenecks such as high index overhead, poor execution of complex joins, and SPARQL query timeouts.

Metabolites, conversions, and catalysis interactions carry no direct species annotation in the data model, since this data is not directly specified by PMN. Species context for these entity types is instead inferred through the graph via a catalysis chain: species X’s enzyme E must be “wp:source” of a “wp:Catalysis” whose “wp:target” is a specific “wp:Conversion” (genes are chained to their enzyme via “wp:TranscriptionTranslation” first). *A. thaliana* (NCBITaxon:3702) is by far the most represented species, with direct annotations on 1,038 of 1,162 pathways (89%), 1,121 gene products (wp:GeneProduct), and 1,325 enzyme nodes (wp:Protein). Via the catalysis chain (the specific “wp:Catalysis” to “wp:Conversion” steps its enzymes actually participate in), *A. thaliana* is linked to 1,354 metabolites (30% of all “wp:Metabolite” nodes), 2,323 biochemical conversions (12% of all “wp:Conversion” nodes), and 4,732 catalysis interactions (38% of all “wp:Catalysis” nodes) (Figure S6).

Biosynthetic gene cluster (BGC) crosslinks, integrated from MIBiG 4.0 and plantiSMASH 2.0 predictions, contribute an additional 7,843 triples to the knowledge graph, including all 43 plant BGCs from MIBiG 4.0, out of which 9 are annotated to *A. thaliana*, and 65 plantiSMASH-detected *A. thaliana* BGCs in the plantiSMASH 2.0 database ^31^. Of these total of 74 *A. thaliana* BGCs, 44 (38 from plantiSMASH and six from MIBiG) share at least one gene (wp:GeneProduct) with a PlantCyc pathway, producing 225 unique (BGC, gene, pathway) crosslinks that involve 66 (BGC, gene) pairs, 58 unique genes (Fig. S7), and 132 unique pathways (Supplementary Table S4; Fig S8; see Data availability). The remaining 30 *A. thaliana* BGCs (27 plantiSMASH and 3 MIBiG), currently have no PlantCyc pathway connection, highlighting potential targets for future pathway annotation across resources and further characterization of clustered regions of biosynthesis into full metabolic pathways. The 3 MIBiG BGCs not connected are those for the biosynthesis of tirucalla (mibig:BGC0001314), retigeranin/arathanatriene (mibig:BGC0001756), and euphol (BGC0002401), fully characterized terpene BGCs with known chemical compound, genes IDs and products, and available literature references. The reason for this missed crosslink is the absence of these pathways from the PlantCyc 17.0 source data, which showcases how the integration of BCGs into a knowledge graph with pathway models can find low-hanging fruits for data curation. The superpathway of phylloquinol biosynthesis (PC721) is the most connected PlantCyc pathway with six BGCs (four plantiSMASH and two MIBiG) through 4 shared genes (Fig S9) out of 23 genes in total (representing 17% of the pathway), showing how independently predicted and curated BGCs can converge on the same PlantCyc pathway.

The resulting current PlantMetWiki RDF knowledge graph is organized as a multi-graph Virtuoso triplestore including seven named graphs (Figure 1.4), each representing a distinct layer of the data model (Supplementary Table S6). The core graph (3,826,567 triples) encodes 2,758 gene products, 3,750 enzymes, 4,577 metabolites, and 54,800 typed interaction edges, of which 19,927 are biochemical conversions and 12,359 are catalytic links. Pathway content is supported by 4,085 distinct literature references annotated with PubMed IDs. A separate properties graph (2,617,839 triples) preserves PlantCyc-specific key–value annotations as RDF triples, making PlantCyc metadata available for SPARQL queries beyond what the core WikiPathways vocabulary captures. This graph represents the basis for future expansions of the current semantic model and supports the use of PlantMetWiki as an expandable framework to support pathway modeling beyond the WikiPathways’ attributes. Dataset provenance, licensing information, versioning metadata, and download locations are described with VoID metadata records following the VoID Schema RDF vocabulary ^32^. These are available as a “void” graph (http://rdf-plantmetwiki.bioinformatics.nl/void) hosted in the Virtuoso database and accessible via the SPARQL endpoint. All workflows used to generate the resource are openly available, and version-controlled, enabling reproducible regeneration and versioning from source data.

### Multispecies pathway modelling enables comparative cross-species pathway analysis and annotation prioritization

A central feature of PlantMetWiki is the representation of pathways as multispecies semantic graphs in which genes, enzymes, reactions, metabolites, and taxonomic annotations can be queried together. This representation enables comparison of pathway coverage across species (e.g., via the previously mentioned catalysis-chain assignment) and supports the identification of annotation gaps that may not be apparent in species-specific pathway views. For each pathway, we classify missing species-reaction annotation as an inferable gap if at least one other species in the same pathway has a direct enzyme annotation for that reaction, and as a non-inferable gap if no species in the pathway does. Inferable gaps mark positions where cross-species evidence can prioritize candidate genes, enzymes, or reactions (wp:Conversions) for further investigation, while non-inferable gaps indicate pathway steps that currently require experimental characterization rather than annotation transfer. To demonstrate this capability, we examined pathways representing distinct biological scenarios, including species-specific specialized metabolism, medicinal alkaloid biosynthesis, and specialized metabolites from non-model plants.

The capsaicin biosynthesis pathway (PWY-5710) illustrates how PlantMetWiki can be used to identify annotation gaps and generate a concrete annotation-transfer hypothesis within an economically important species and well-characterized metabolite. PlantMetWiki contains annotations for four Capsicum taxa: three species (*Capsicum annuum, Capsicum chinense*, and *Capsicum frutescens*) and the genus-level node Capsicum. *C. annuum* is the most extensively represented of these taxa across the entire PlantMetWiki core graph, with direct enzyme annotations in 34 pathways. However, querying PWY-5710 reveals an annotation gap for this species, with *C. annuum* showing no directly annotated gene or enzyme nodes to the capsaicin biosynthesis pathway. This absence reflects a limitation of the source curation and demonstrates how cross-species querying can reveal pathway-specific gaps that are difficult to identify through conventional pathway browsing. In total, nine species are represented in the pathway, including *C. chinense, C. frutescens*, and four phylogenetically distant species that contribute annotations to individual pathway steps. The pathway comprises seven biochemical conversions (wp:Conversion steps; GPML diagram branch-point duplicates excluded), of which only 10 of the 63 possible species–reaction combinations (15.9%) are directly annotated across all nine species combined (Fig. S10-12). Capsaicin synthase (MONOMER-13528, UniProt O81646; Figure 3), the pathway’s terminal acyltransferase, is annotated in *C. chinense* and catalyzes a reaction (RXN-8925) that is unannotated in *C. annuum*, *C. frutescens*, and the genus-level *Capsicum* node. The *C. chinense* capsaicin synthase sequence is therefore a direct, named candidate for targeted annotation of the missing *C. annuum* step, showing how PlantMetWiki’s reaction-level resolution converts a generic annotation gap into a specific, testable hypothesis. More broadly, by exposing which reactions lack species-specific evidence and identifying related taxa to support annotation transfer, PlantMetWiki enables systematic prioritization of candidate pathway annotations for *Capsicum* species.

**Figure 3:**
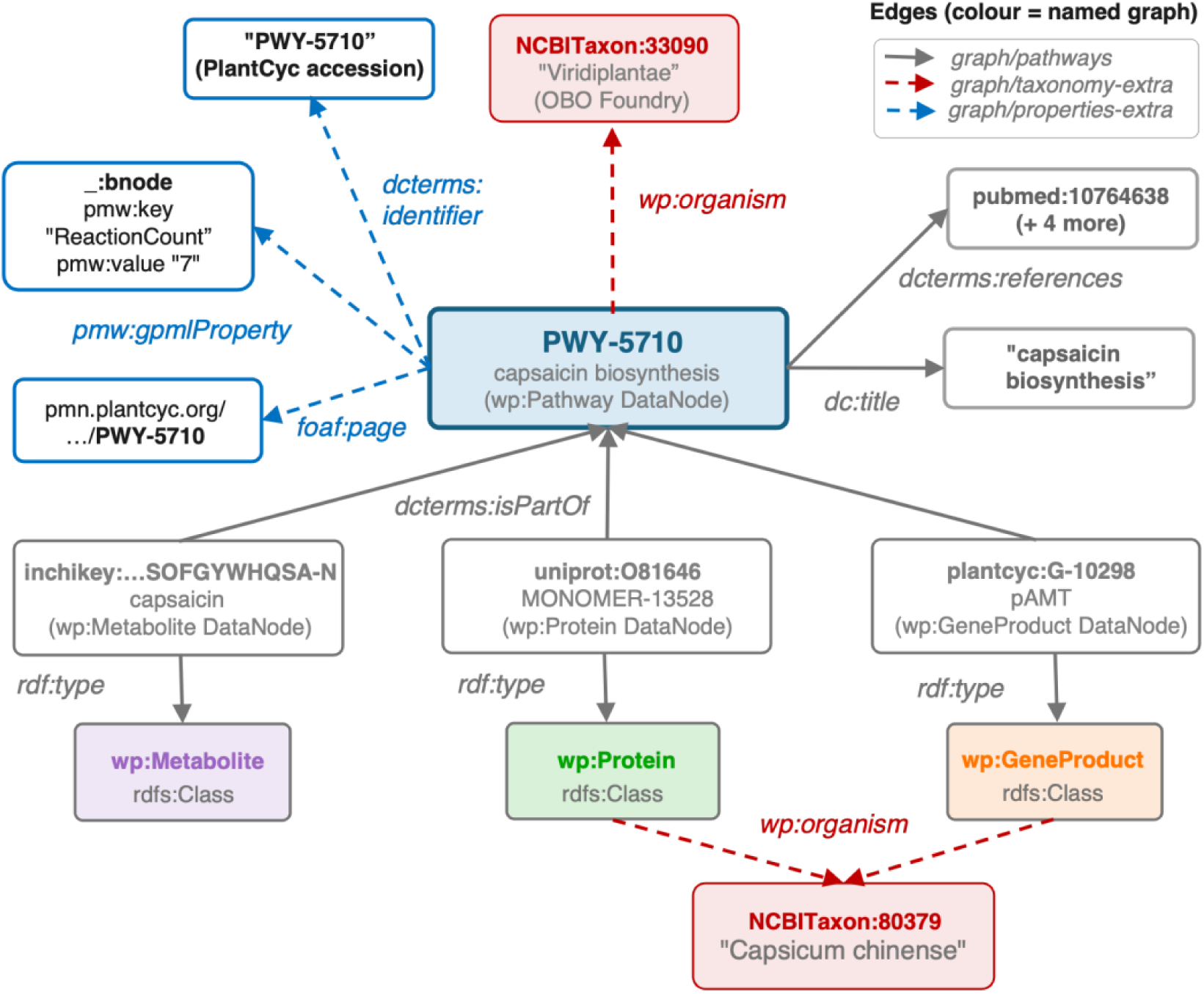
Semantic model of the capsaicin biosynthesis pathway (PWY-5710) across named graphs. Capsaicin biosynthesis (PWY-5710, “wp:Pathway”) and DataNode MONOMER-13528 (*Capsicum chinense* capsaicin synthase, UniProt O81646, “wp:Protein”) shown across three named graphs. Solid grey edges originate from graph/pathways (core WikiPathways RDF, generated by gpml2rdf); dashed red edges from graph/gpml-taxonomy-extra (species enrichment layer with NCBI Taxonomy); dashed blue edges from graph/gpml-properties-extra (PlantCyc property layer). The core “wp:organism” on PWY-5710 is assigned in the RDF generation with the taxonomy-extra layer and mapped to the OBO Foundry IRI graph, resulting in NCBITaxon:33090 (Viridiplantae) mapped to the pathway and NCBITaxon:80379 – among others – mapped to the individual Protein and GeneProduct nodes. The Metabolite, Protein and GeneProduct Datanodes are connected to PWY-5710 via “dcterms:isPartOf” in graph/pathways.

The hyoscyamine and scopolamine biosynthesis pathway (PWY-7341) illustrates a complementary scenario: unlike capsaicin, where a single congeneric species supplies one direct candidate for one missing reaction, here the missing annotation of a single target species must be reconstructed from several partial, non-redundant sources. *Atropa belladonna* (NCBITaxon:33113) is a plant from the Solanaceae family and a hyoscyamine-rich producer of tropane alkaloids from which atropine – the racemic mixture formed from its principal alkaloid, L-hyoscyamine – takes its name ^32^. *A. belladonna* contains only a single directly annotated enzyme, covering just one of the pathway’s 16 biochemical conversions, despite 13 Solanaceae species being annotated in the same pathway. *Hyoscyamus niger* (NCBITaxon:4079), by comparison, contains four annotated enzymes mapping to two of the 16 reactions, and *Datura stramonium* (NCBITaxon:4076) contributes a further three reactions, together providing five reaction-level candidates (none overlapping with *A. belladonna*’s single annotated step), for completing the *A. belladonna* pathway annotation. Across the pathway, only 16 of 208 possible species–reaction combinations are directly annotated (7.7%), indicating substantial annotation sparsity. No single relative can resolve this gap on its own: the annotated Solanaceae species each contribute to non-overlapping reaction-level coverage of biosynthetic knowledge: reconstructing the biosynthetic routes for hyoscyamine and scopolamine production and identifying candidate enzymes and reactions for comparative analysis in *A. belladonna* requires tapping into this pool (Figure S13). Notably, this transfer strategy mirrors an experimentally validated precedent: heterologous expression of the *H. niger* H6H gene in *A. belladonna* redirects alkaloid flux almost entirely toward scopolamine, confirming that a computationally prioritized donor species can correspond to a biologically validated gene transfer supporting plant metabolic engineering. This example illustrates how PlantMetWiki can integrate fragmented biosynthetic knowledge distributed across many related species into a single queryable evidence base, supporting comparative exploration and hypothesis generation of candidate pathway components, as well as metabolic engineering targets.

Avenacin A-1 ^33^ biosynthesis (PWY-7476) illustrates the limits of enzyme-count-based cross-species comparison. Avenacin A-1 is an antifungal triterpenoid saponin produced by *Avena strigosa* (NCBITaxon:38783). *A. thaliana*, which lacks the oat-specific avenacin pathway ^25^, contains annotations for seven enzymes to reactions in the same pathway, corresponding to 87.5% of the eight enzyme-associated bioconversion (wp:Conversion) steps annotated in the native producer. At the reaction level, however, this apparent overlap does not hold: of the pathway 13 biochemical conversions, *A. strigosa* covers nine, while *A. thaliana’*s seven annotated enzymes collectively map onto only two reactions, both already covered in *A. strigosa* itself. This is because PlantMetWiki’s species-reaction mapping reflects which “wp:Conversion” node an enzyme is linked to via “wp:Catalysis” (i.e., shared position in the pathway graph). *A. thaliana’*s seven enzymes are multiple paralogs independently annotated to the same two broadly conserved, upstream triterpenoid-backbone reactions shared across many species, rather than to the seven downstream, avenacin-specific tailoring reactions unique to Avena ^25^ (Figure S14)*. A. thaliana* therefore provides no reaction-level evidence that is not already present in the native producer, despite its near-equivalent enzyme count. The remaining four reactions that have no “wp:Catalysis” evidence from any of the 15 species annotated in this pathway, including *A. strigosa* itself, constitute a non-inferable gap: PlantMetWiki currently provides no cross-species evidence that could support annotation of these steps, highlighting them as priorities for experimental characterization rather than annotation transfer (Figure S14).

Monolignol biosynthesis (PWY-361) demonstrates the value of PlantMetWiki for breadth-based, family-level inference across an agronomically critical pathway. With 29 annotated species spanning model plants, major crop grasses, trees, and non-model taxa, PWY-361 is the most taxonomically diverse specialized pathway in PlantMetWiki. *A. thaliana* provides the most complete coverage, with 19 annotated enzymes spanning all 15 of the pathway’s 15 biochemical conversions (100%), while 28 of the 29 species cover fewer than 40% of this enzyme depth, including all economically important Poaceae crops present in the graph: rice, maize, sorghum, switchgrass, ryegrass, and wheat. Querying the reaction coverage matrix reveals that seven of these 15 reactions are annotated in *A. thaliana* but in no grass species, representing the highest-priority inference targets for lignin pathway annotation in cereals and bioenergy crops. Within the Poaceae, the combined annotation footprint of all grass species together covers the remaining ing eight of 15 reactions (all of which are also covered in *A. thaliana*), providing within-family transfer candidates that are phylogenetically closer to the target crops than the *A. thaliana* reference. Improving monolignol pathway annotation in these species has direct relevance for the rational design of modified-lignin varieties for cellulosic bioenergy and improved biomass digestibility (Figure S15).

These four scenarios illustrate different uses of the cross-species pathway representations in PlantMetWiki: hypothesis generation for annotation transfer of reaction steps and related enzymes from a related species (capsaicin), pooled reconstruction of bioconversion steps from multiple complementary sources (hyoscyamine/scopolamine), difference between cross-species coverage of precursor and tailoring enzymatic steps (avenacin), and breath-based inference within a large, taxonomically diverse family (monolignol). By combining pathway structure with species annotations in a common semantic framework, PlantMetWiki enables researchers to identify annotation gaps, retrieve candidate annotations from related species, and prioritize biosynthetic hypotheses for experimental follow-up. The public SPARQL endpoint provides reproducible access to these analyses, and all queries used in this study are available through the accompanying notebooks and example queries (see Data availability section).

Applying this reaction completeness quantification strategy across PlantMetWiki, reveals the proportion of pathway–species annotation gaps for which supporting evidence is available elsewhere in the knowledge graph. Across 1,093 pathways with at least one species-tagged (wp:organism) enzyme annotation, the complete reaction-by-species matrix comprised 29,061 possible species–reaction combinations. Of these, 7,012 (24.1%) contained direct annotations, while 13,922 (47.9%) represented inferable gaps supported by annotations in at least one other species (Figure 4A). The remaining 8,127 combinations (28.0%) corresponded to reactions lacking evidence across all species represented in the pathway (Figure 4). These results indicate that almost two thirds (63.1%) of all currently unannotated pathway positions in PlantMetWiki are potentially addressable through cross-species evidence transfer. The remaining 36.9% reflect pathway steps for which no evidence is currently available anywhere in the knowledge graph and therefore require additional experimental characterization rather than annotation propagation. The pathway case studies presented above showed consistently higher levels of inferable annotation space than the global average, as reflected in Figure 4B. Capsaicin biosynthesis contained 55.6% inferable gaps, hyoscyamine and scopolamine biosynthesis 48.6%, avenacin A biosynthesis 56.9%, and monolignol biosynthesis 83.7%, compared to 47.9% globally. This reflects the selection criteria used to identify these case studies (≥3 annotated species, ≥8 biochemical conversions, ≥6 enzyme annotations), which favor pathways with sufficient cross-species annotation density for inferable gaps.

**Figure 4:**
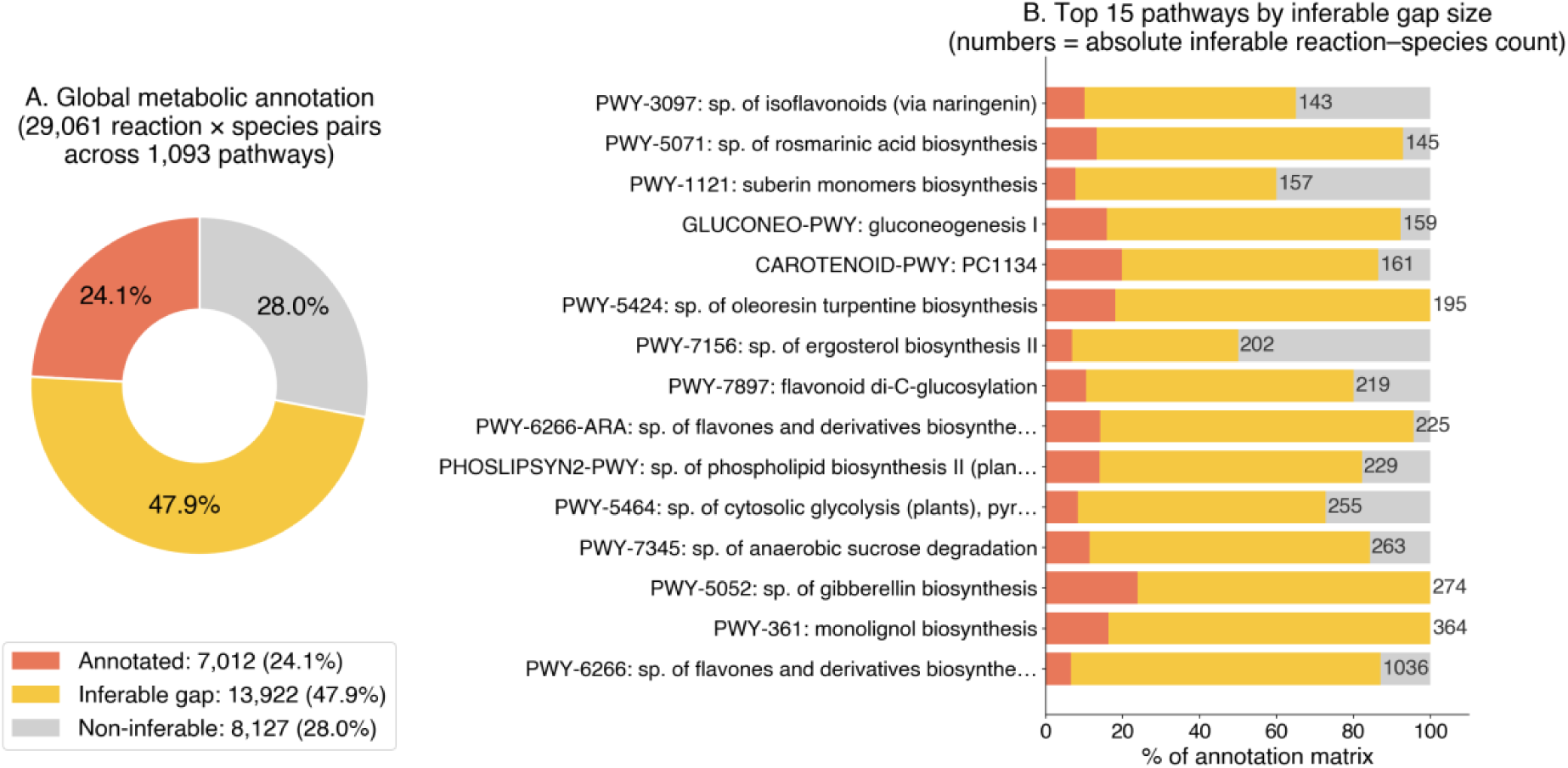
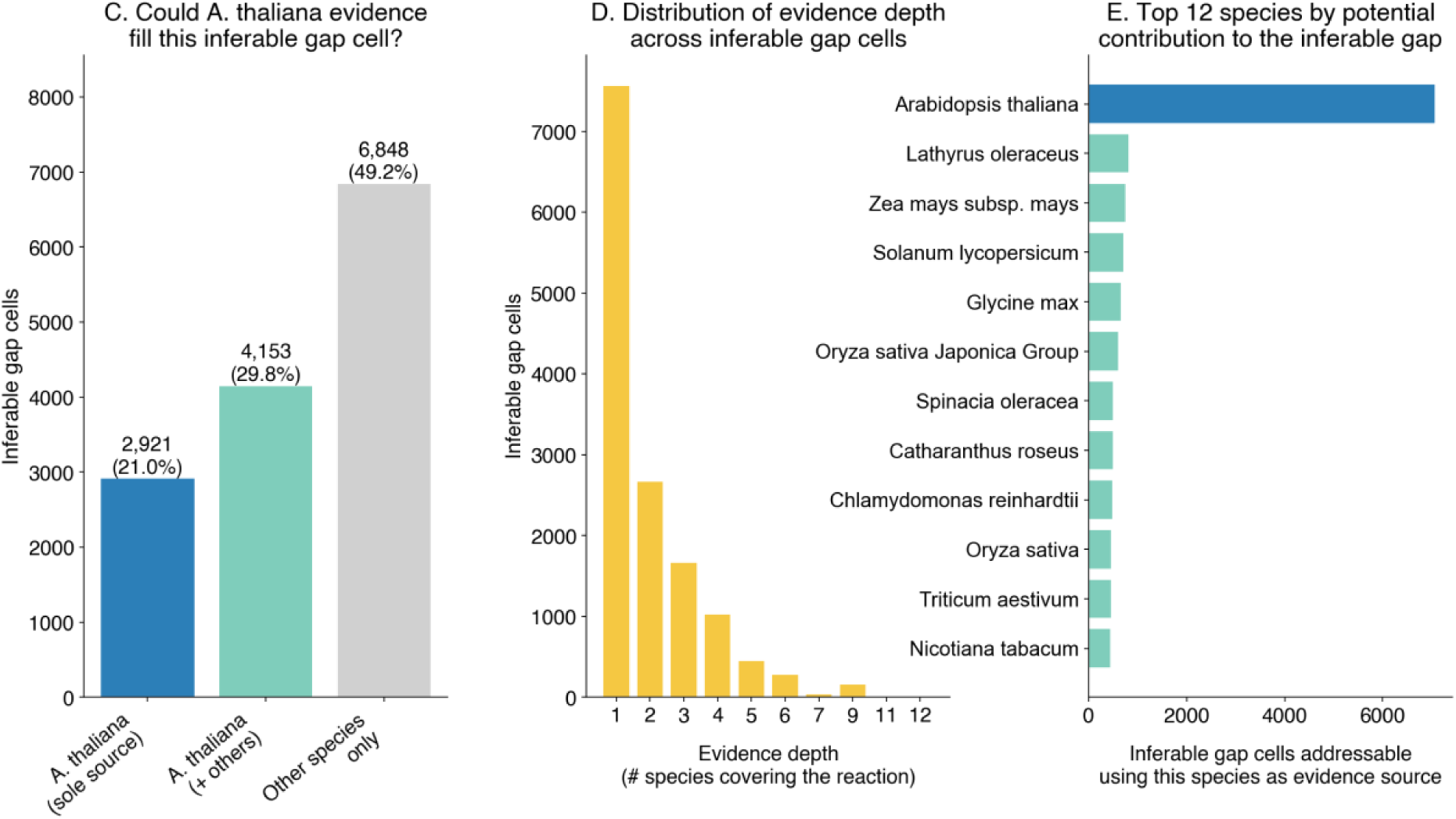
Metabolic annotation matrix and cross-species inference potential of PlantMetWiki and contribution of *A. thaliana*. (A) Global breakdown of all (reaction × species) pairs across all pathways with ≥1 species annotation. "Annotated" = “wp:Catalysis” link exists for the species–reaction pair; "Inferable gap" = gap where ≥1 other species in the same pathway has a Catalysis link for the same reaction (cross-species transfer candidate); "Non-inferable" = reaction completely uncharacterized across all species in the pathway; “sp” = super pathway. (B) Top 15 pathways ranked by absolute inferable gap (number of reaction–species pairs addressable by cross-species transfer), with color breakdown showing the proportion of each category. Monolignol biosynthesis is the 14^th^ pathway in the rank. (C) Breakdown of the 13,922 inferable gap cells by whether *A. thaliana* provides evidence (sole source; one of several sources; no A. thaliana evidence). (D) Distribution of evidence depth (number of species with enzymes annotated to a given reaction) across inferable gap cells. (E) Top species ranked by total inferable gap cells could help resolve.

While the contribution of the model plant *A. thaliana (*the most extensively annotated species in PlantMetWiki) is large, the inferable annotation gaps rely on the broader diversity represented in PlantMetWiki. Of the 13,922 inferable gap cells (species-reaction pair with no direct enzyme annotation for that species), 2,921 (21.0%) could only be resolved using *A. thaliana* annotations (i.e., *A. thaliana* is the sole species covering that reaction within its pathway), a further 4,153 (29.8%) have *A. thaliana* as one of several possible sources, and the remaining 6,848 (49.2%) rely entirely on evidence from other species. Overall, *A. thaliana* annotations could contribute to approximately half (50.8%) of the inferable gap, while the other half is independent of *A. thaliana* and draws on a long tail of species, most prominently *Lathyrus oleraceus, Zea mays, Solanum lycopersicum, Glycine max,* and *Oryza sativa* (Figure 4). In terms of evidence robustness, 54.4% of inferable gap cells have only a single potential source species, while the remaining 45.6% are supported by two or more independent species, providing mutually corroborating evidence. These results indicate that, while *A. thaliana*’s annotation depth makes it the single most valuable reference species in PlantMetWiki, cross-species inference in this resource is not solely dependent on a model-plant reference and draws on a broad repertoire of annotated taxa.

### Connecting plant metabolism to the Linked Open Data ecosystem through mapping and federation

The PlantMetWiki SPARQL endpoint positions the resource as a key tool in the LOD ecosystem through two complementary strategies: direct cross-resource identifier mapping (via BridgeDB), which materializes links to external chemical databases inside PlantMetWiki knowledge graph, and federated queries, which reach any LOD resource at query time through shared identifiers, such as InChIKeys ^33^. Together, these allow PlantMetWiki metabolites to be queried alongside resources such as Wikidata, ChEBI ^34^, and PubChem ^35^ within a single SPARQL query. This interoperability addresses a long-standing disconnect in plant biochemistry: pathway databases capture how and in which species metabolites are produced, whereas chemical databases focus on molecular properties, biological activities, toxicology, and applications. In Wikidata alone, the production of atropine (Q26272) is not connected to the *A. belladonna* plant (Q156091): mapping and federation bridge these complementary resources.

Nearly 90% of PlantMetWiki metabolites can be linked to Wikidata entries via federated queries using their InChIKey, which Wikidata uses as a standard chemical identifier (wdt:P235). Of the 4,577 metabolite nodes in the resource, 4,111 (89.8%) carry an InChIKey and can be joined to Wikidata within a single query. Additional federation points are provided through ChEBI (174), MetaNetX (172), PubChem Compound (75), and KEGG Compound (43) metabolite identifiers (Figure S18). However, most source metabolites carry only a single primary identifier from these resources and almost no cross-references between chemical databases, so reaching ChEBI, PubChem, HMDB or Wikidata for a given compound still requires a live federated lookup.

To make these links first-class and directly queryable, we integrated the BridgeDb metabolite ID-mapping database (build 20260102; derived from HMDB, ChEBI, and Wikidata) into the GPML-to-RDF conversion. This materialized cross-references for 3,198 of 4,577 metabolites (69.9%), stored under dedicated “”wp:bdb* predicates: 3,059 Wikidata, 2,926 PubChem, 2,805 ChEBI, 2,317 ChemSpider, 1,120 KEGG, 975 HMDB, and 510 LipidMaps links (Figure 5). BridgeDb thus densifies the sparse PlantCyc-derived identifiers into multi-database hubs, raising ChEBI coverage from 174 to 2,805 metabolites, PubChem from 75 to 2,926, and KEGG from 43 to 1,120. BridgeDB further adds Wikidata, ChemSpider, HMDB, and LipidMaps links, which are otherwise absent from the source PlantCyc altogether. This results in the majority of mapped metabolites now linking to four or more databases. The two strategies are therefore complementary: whereas federation reaches a Wikidata entry at query time by matching on the InChIKey, BridgeDb stores the other database entities as an explicit link on the metabolite node, so the connection is resolved once, offline, and directly traversable. InChIKey-based federation remains available for the 26.4% of metabolites not covered by BridgeDb.

**Figure 5:**
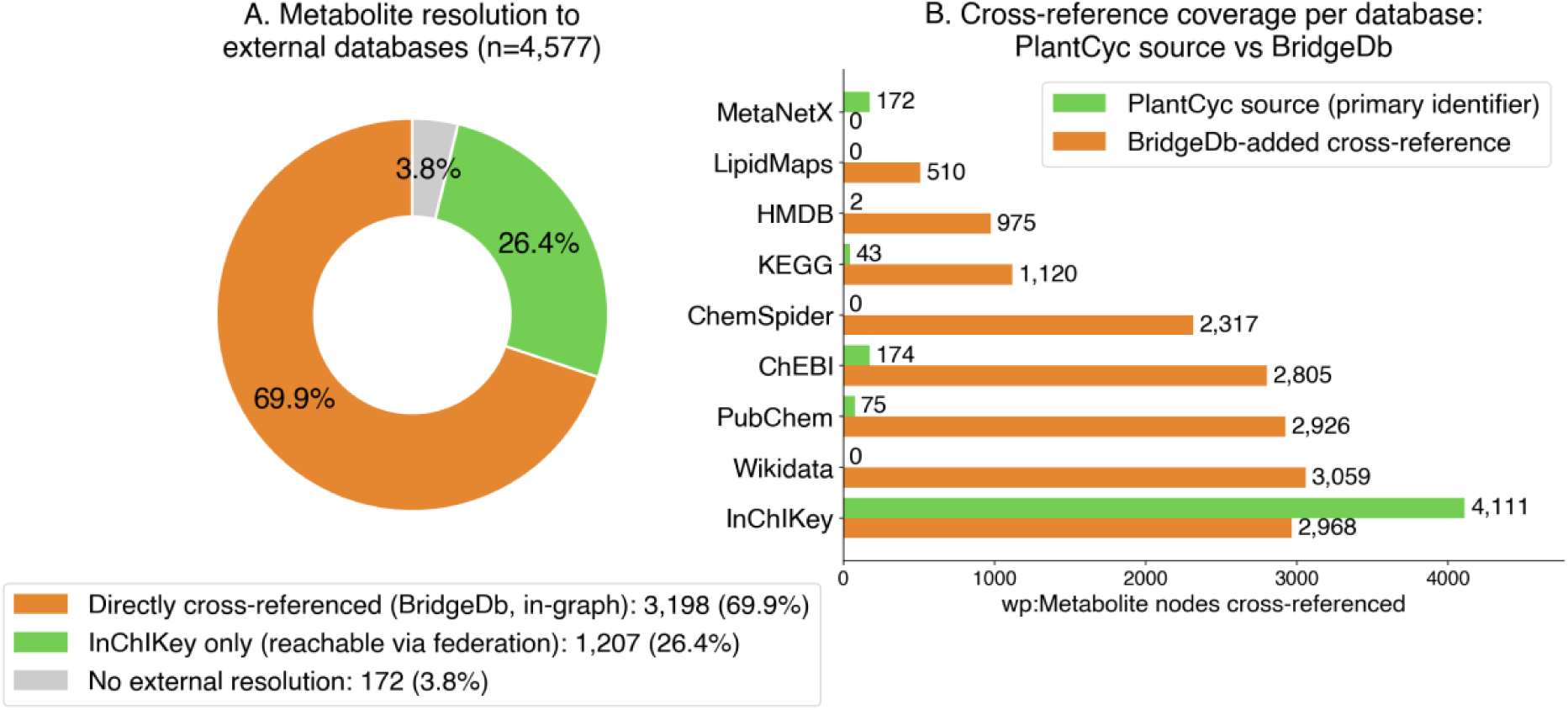
Metabolite cross-reference coverage: direct BridgeDb mapping complements InChIKey federation. (A) How the 4,577 wp:Metabolite nodes resolve to external chemical databases: 3,198 (69.9%) carry ≥1 BridgeDb-materialised cross-reference and are resolvable directly in the graph without federation; 1,207 (26.4%) carry only an InChIKey and remain reachable through query-time federation to Wikidata (wdt:P235) and other InChIKey-indexed resources; 172 (3.8%) have neither. (B) Cross-reference coverage per database, comparing the single PlantCyc source identifier each metabolite carries with the BridgeDb-added wp:bdb* cross-references. PlantCyc supplies mostly a lone InChIKey with few direct database links (ChEBI 174, PubChem 75, KEGG 43); BridgeDb (build 20260102: HMDB, ChEBI, Wikidata) raises these to 2,805 ChEBI, 2,926 PubChem and 1,120 KEGG and adds Wikidata (3,059), ChemSpider (2,317), HMDB (975) and LipidMaps (510) links absent from the source, so most mapped metabolites become hubs to ≥4 databases. MetaNetX is not part of the BridgeDb metabolite build and remains source-only. Mappings are added under separate predicates and never overwrite the curated identifier.

Because BridgeDb writes these cross-references under separate predicates, storing the external links directly on each metabolite, it augments rather than overwrites the curated PlantCyc identifier: the BridgeDb-derived InChIKey agreed exactly with the source InChIKey for 2,904 of the 2,904 metabolites carrying both (100%). Only four metabolites received a BridgeDb InChIKey sharing the 2D skeleton but differing in the stereochemistry layer: maltose, hemanthamine, lanatoside C, and polyneuridine aldehyde. Three cofactors (FMN, protoheme, and molecular hydrogen, H₂) received structurally unrelated key values, reflecting the well-known instability of InChIKeys for charged and metal-coordinated species rather than mapping errors. At 0.14% stereo-variants (4/2,904), source-to-source agreement is near-total, so the mapping is highly reliable. The rare stereochemical discrepancies mirror the difference in stereochemical completeness between a pathway-centric source (PlantCyc) and compound-centric ones (PubChem/Wikidata, which specify the full stereochemistry configuration). Exact-stereo queries can always rely on the curated source "dcterms:identifier”, reinforcing the value for cross-database quality control.

Two case studies illustrate complementary directions enabled by BridgeDB mapping and federation: enriching PlantMetWiki with external chemical and pharmacological annotation (alpha-tomatine) and using external knowledge to expose gaps within PlantMetWiki itself (theobromine). Alpha-tomatine, a steroidal glycoalkaloid produced by *Solanum lycopersicum* (tomato) and related Solanaceae species, is directly linked to Wikidata (Q288051), PubChem CID (28523), and ChEBI (9630) via BridgeDB. Federation retrieves annotations from Wikidata describing alpha-tomatine as an antifungal agent, anti-infective agent, and bitter compound, and from PubChem adding toxicological studies and records of industrial applications in cosmetics and as an agricultural fungicide. This information is now available on the PlantMetWiki UI thanks to queries targeting both resources. Additionally, PlantMetWiki supplies what neither resource provides alone: the three distinct glycoforms of tomatine (alpha, beta-1, and gamma), each represented as a separate metabolite node with an InChIKey identifier, placed in their biosynthetic context within the glycoalkaloid metabolism superpathway (PC426) and the dedicated alpha-tomatine biosynthesis pathway (PC1159), with species-level annotation spanning five *Solanum* spp. A single federated query can therefore traverse from the biological activity recorded in Wikidata, through the biosynthetic route encoded in PlantMetWiki, to the specific plant species and enzymatic steps responsible: a workflow that would otherwise require sequential manual lookup across three separate databases.

Theobromine illustrates federation working in both directions simultaneously. Theobromine is a well-characterized methylxanthine alkaloid (Wikidata Q206844, ChEBI 28946, CAS 83-67-0) with extensive pharmacological annotations documented across these resources, including its roles as a bronchodilator, vasodilator, and nootropic. PlantMetWiki adds pathway-level context: theobromine participates in seven pathways, spanning two dedicated biosynthesis routes resulting from convergent evolution of methylxanthine metabolism in plants ^35^, with enzymatic annotations linking *Coffea arabica* (12 annotated gene and enzyme nodes) and *Camellia sinensis* (4 nodes) as the primary gene-annotated species (Fig S16-17). The resource also captures the metabolic neighborhood of theobromine within the methylxanthine family, with caffeine, theophylline, and 7-methylxanthine represented as co-metabolites across overlapping pathways, providing a graph-queryable view of the xanthine alkaloid network that Wikidata and ChEBI currently do not represent at the pathway level.

The theobromine example drives federation in the reverse direction, exposing a significant gap in PlantMetWiki itself. Wikidata records *Theobroma cacao* (cocoa) - the plant that gives theobromine its name – as a confirmed theobromine-producing taxon via the found-in-taxon property (wdt:P703), alongside *Coffea arabica*, *Camellia sinensis*, *Cola nitida*, and *Paullinia cupana*. Yet, the PlantCyc source data and PlantMetWiki core graph contain no annotated gene or enzyme nodes for *T. cacao* in any of the theobromine biosynthesis pathways, despite it being the historically and commercially primary producer of the compound. This gap is invisible when exploring PlantCyc alone but becomes immediately apparent once the theobromine InChIKey is joined in a federated query. The finding identifies *T. cacao* theobromine pathway annotation as a high-priority curation target and shows how linked data can function as an integration framework and as a mechanism for quality control and gap detection. Beyond these individual compounds, InChIKeys help us quantify how much of PlantMetWiki’s chemistry is absent from external resources. A complete federated scan of all 4,111 InChIKey-annotated PlantMetWiki metabolites against Wikidata showed that 2,876 metabolites (70.0%) currently have an exact InChIKey match in Wikidata, whereas 1,235 (30.0%) are absent. These include compounds such as the acylated sucrose ester 4-(3-methylbutanoyl)sucrose (XUZMIKFKZAUSGG-IVHWZWKWSA-N), the acylated flavonoid glycoside isorhamnetin 3-O-(3″,6″-di-O-4-coumaroyl)glucoside (CQKFGYQBBQNDQG-NKXGADSASA-N), and the *Allium* organosulfur volatile (Z)-butanethial oxide (BQXJLKVZTDDERJ-UHFFFAOYSA-N), which are present in PlantMetWiki with full pathway context but have no corresponding Wikidata entry.

Most of these absences reflect alternative molecular representations rather than genuinely missing chemistry. To separate genuine knowledge gaps from stereoisomers, tautomers, glycoforms, or salt forms of already represented molecules, we compared the unmatched metabolites at the InChIKey connectivity layer, represented by the first 14-character block, which encodes the molecular skeleton independent of stereochemistry, isotopic composition, or protonation state. Of the 1,235 unmatched metabolites, 814 (65.9%) share a skeleton with at least one Wikidata compound (Figure 6A,C), therefore representing relatively soft knowledge gaps. For instance, PlantMetWiki momilactone A record (InChIKey MPHXYQVSOFGNEN-QCZCYNOHSA-N) carries no exact Wikidata match, yet shares its skeleton with four Wikidata entries, including the Wikidata entry for momilactone A itself (Q27104650), which carries a different InChIKey (MPHXYQVSOFGNEN-JGHPTVLTSA-N; PubChem CID 162644). The two representations are the same molecule (identical formula, C₂₀H₂₆O₃, and identical atomic connectivity) and differ only in the InChI stereochemistry layer: three stereocentres (C13, C14, C16) are left undefined in the PlantMetWiki/PlantCyc record but are fully specified in the Wikidata/PubChem record (Supplementary Table S7). The apparent absence is, therefore, not missing chemistry but a difference in the completeness of stereochemical annotation between a pathway-centric source (PlantCyc) and a compound-centric one (PubChem/Wikidata). The remaining 421 metabolites (34.1% of the unmatched set; 10.2% of all InChIKey-annotated metabolites) have no skeleton-level match, indicating that Wikidata contains no structurally related representation and marking them as the strongest candidates for genuinely novel contributions to the LOD ecosystem (Figures 6A, C; Supplementary Table S7).

**Figure 6:**
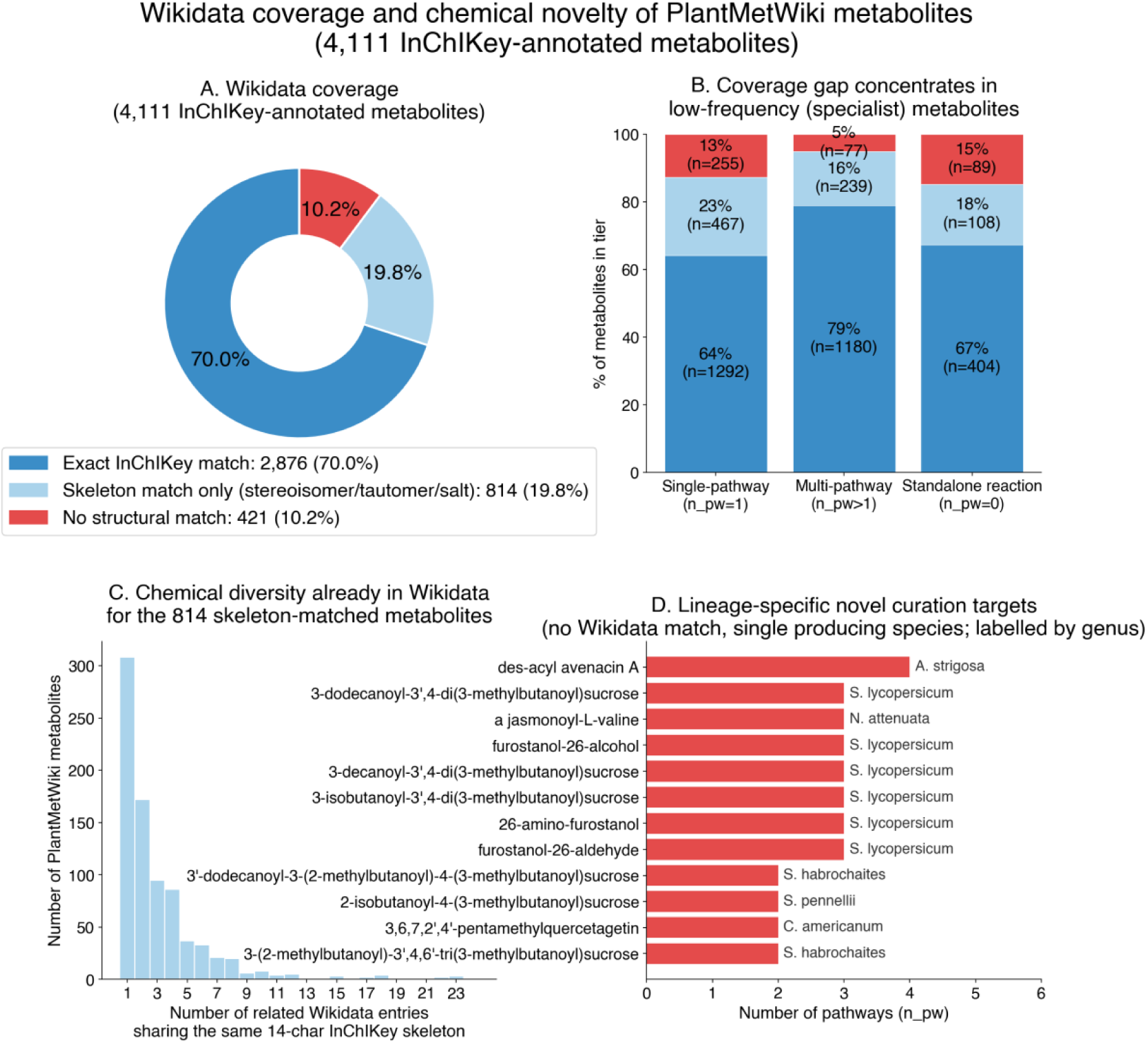
Wikidata coverage and chemical novelty of the 4,111 InChIKey-annotated metabolites. (A) Overall split into exact InChIKey matches (blue, already in Wikidata), skeleton-only matches (light blue, Wikidata holds a stereoisomer/tautomer/salt form of the same molecule, but not this exact form), and no structural match (red, the molecule is entirely absent from Wikidata). (B) The same three categories, broken down by how often each metabolite appears across pathways: single-pathway (n_pw=1, most specialist), multi-pathway (n_pw>1), and metabolites belonging to standalone reactions (no membership to “PC*” pathways), such as 8’-apo-β-carotenal, 1-heptanal, trans-4-hydroxy-L-proline, peroxynitrite, methyl-β-D-galactoside. (C) For the 814 skeleton-matched metabolites, the distribution of how many *other related entries (different stereoisomers/forms of the same skeleton) Wikidata already holds: most skeletons have only 1–2 related Wikidata entries, but a few (e.g., kaempferol glycosides) already have many. (D) The 12 highest-priority lineage-specific novel-scaffold metabolites (compounds absent from Wikidata (no skeleton match) that are produced by exactly one annotated species) ranked by pathway count and labelled with the producing genus. Restricting to single-species compounds excludes reaction intermediates with no organism-annotated enzyme and highlights the taxonomically restricted specialized metabolites PlantMetWiki uniquely holds. The full ranked list of all 421 novel candidates is in Supplementary Table S7.

Specialized chemistry drives the Wikidata annotation gap, evident in both pathway distribution and, more strongly, the taxonomic distribution of the absent metabolites. Absent metabolites concentrate among compounds occupying few pathways: single-pathway metabolites (n_pw = 1) and metabolites present only in standalone reactions (n_pw=0) are absent from Wikidata at 35.8% (722/2,014) and 32.8% (197/601), respectively, versus 21.1% (316/1,496) for multi-pathway metabolites (Figure 6B). The taxonomic distribution showed the same trend more sharply. Following the catalysis chain (wp:Metabolite → wp:Conversion → wp:Catalysis → wp:Protein → wp:organism), Wikidata absence declines steeply and monotonically as more species are known to make a compound: from 37.8% for metabolites that no annotated species produces (612/1,617), to 30.6% for those made by a single species (476/1,554), 18.7% by two or three species (124/664), and below 10% for those synthesized by four or more (8.5%, 20/235 at four-nine species; and 7.3%, 3/41 at ten or more) (Figure S19).

Single-pathway metabolites are overwhelmingly single-species (86% are made by ≤1 annotated species), and 93% of the 421 novel-scaffold compounds are produced by at most one annotated species, consistent with the highly specialized nature of plant metabolism. The novel-scaffold set (no structurally related entries in Wikidata) is dominated by taxonomically restricted metabolites (compounds present in a single PlantMetWiki pathway and produced by a single annotated species). These span diverse classes of specialized metabolism: the triterpenoid-saponin precursor des-acyl avenacin A (WDHZXDLRQGPOGR-WGJSNZCWSA-N; *Avena strigosa*, four pathways), the jasmonate conjugate jasmonoyl-L-valine (*Nicotiana attenuata*, three pathways), and the polymethylated flavonoid 3,6,7,2′,4′-pentamethylquercetagetin (*Chrysosplenium americanum*, two pathways), each confined to a single producing species yet with no structural counterpart in Wikidata (Figure 6D; Supplementary Table S7). This finding establishes PlantMetWiki as a resource rich in lineage-specific plant specialized compounds.

These results demonstrate that the relationship between PlantMetWiki and Wikidata is bidirectional. Wikidata enriches PlantMetWiki with biological-role, taxonomic, pharmacological, and toxicological information, while PlantMetWiki contributes pathway context and at least 421 largely specialized metabolites for which no structurally related representation currently exists in Wikidata. Beyond identifying gaps, federation also exposes cross-resource curation differences: as shown for momilactone A, exact-match absences can reflect differing stereochemical completeness rather than missing chemistry, so the two-tier (exact plus skeleton-level) comparison both separates genuine novelty from representational variants and flags concrete reconciliation targets. Federated querying thereby provides a quantitative framework for assessing knowledge completeness across biological knowledge graphs, for detecting curation discrepancies between them, and for prioritizing future curation where it is most needed. The SPARQL endpoint and federated query patterns demonstrated in the accompanying notebook (see Data availability section) provide the infrastructure to systematically contribute this pathway context to Wikidata as a community curation effort.

## Discussion

### PlantMetWiki enables FAIR multispecies representation of plant metabolism for cross species comparison, pathway gap analysis, and candidate finding

PlantMetWiki transforms species-centric pathway information into a FAIR multispecies knowledge graph that exposes previously hidden relationships and annotation gaps across pathway, genomic, taxonomic, and chemical resources. While Plant Metabolic Network remains the authoritative source of pathway curation, PlantMetWiki provides a complementary semantic representation optimized for interoperability, integration, and reuse, using standard RDF vocabularies, persistent identifiers, machine-readable provenance, and SPARQL-based access. This LOD representation enables pathway knowledge to be queried together with external biological, chemical, and genomic resources without requiring duplication of those resources or manual data integration.

Semantic integration via existing vocabularies, such as the WikiPathways (wp:) one, functions not only as a mechanism for interoperability but also as a framework for identifying knowledge gaps across resources. Cross-species pathway queries revealed pathway positions lacking annotation in individual species despite supporting evidence elsewhere in the graph. Integration with biosynthetic gene cluster resources exposed clustered genomic regions that currently lack pathway assignments. Federated querying against Wikidata revealed both representation-level discrepancies arising from alternative stereochemical forms and 421 metabolites for which no structurally related representation currently exists. These examples demonstrate that incompleteness can become visible only when complementary knowledge sources are considered simultaneously. The distinction between exact InChIKey matches and skeleton-level matches illustrates the importance of chemical normalization when assessing interoperability between biological knowledge graphs, as many apparent absences reflect alternative stereochemical or protonation representations rather than genuinely missing chemistry. The reliable determination of stereochemistry and the subsequent reporting of certainty and uncertainty of chiral centers in compounds are challenging aspects that LOD and federation can help to normalize across resources with different curation depth. As stereochemistry plays a key role in specialized compound bioactivity, with increasing reporting standards and uptake ^36^, resources like PlantMetWiki can help to uncover specific stereochemical patterns across multiple species.

Integration of biosynthetic gene cluster annotations illustrates how semantic representations can connect traditionally separate views of plant metabolism. Pathway databases describe biochemical transformations, whereas genome-mining resources focus on clustered genomic loci. By linking PlantCyc-derived pathways to experimentally validated clusters from MIBiG and predicted clusters from plantiSMASH, PlantMetWiki enables these complementary pathway representations and collections to be explored jointly. The resulting crosslinks connect clustered and unclustered biosynthesis within a common framework and provide a basis for investigating how genomic organization relates to metabolic pathway structure. This demonstrates how FAIR knowledge graph approaches and Semantic Web technologies can bridge resources that were originally developed for different biological questions or with different search methods (such as genome mining in the case of plantiSMASH and MIBiG). Thanks to these crosslinks, the PlantCyc pathways can now be connected to clustered regions in the genome, which were not available either via PMN or MIBiG repositories. By linking clustered and unclustered biosynthesis, PlantMetWiki will further help to understand the complex metabolic network in plants, and their chromosomal arrangement.

A key advantage of PlantMetWiki is its ability to connect pathway knowledge with biosynthetic gene cluster (BGC) information, bridging curated metabolic pathways with genome mining approaches, and fill the knowledge gaps in complex and partially clustered biosynthesis. By integrating experimentally validated clusters from MIBiG 4.0 and predicted clusters from plantiSMASH 2.0, the resource enables users to explore relationships between clustered and unclustered biosynthetic routes and to identify candidate genes associated with specific metabolic conversions and fills annotation gaps across resources focusing on clustered or unclustered pathways. For instance, the alpha-tomatine BGC has been key to characterize two related glycoalkaloid biosynthetic pathways involving orthologous genes between tomato (*Solanum lycopersicum*) and potato (*Solanum tuberosum*) and spanning across two biosynthetic loci: a cluster on chromosome 7 and an additional duplicated region on chromosome 12 ^23^. Because of this incomplete clustering, alpha-tomatine was retired from the MIBiG 4.0 update, resulting in no available mapping of these loci to plantiSMASH 2.0 output. The plantiSMASH clusterBLAST module reports the similarity of *S. lycopersicum* BGCs with ten other BGCs across species, to facilitate comparative analysis. By mapping plantiSMASH-detected regions to PlantCyc pathway annotations, PlantMetWiki facilitates further comparative studies involving pathways across species, resources, and genomic context.

PlantMetWiki provides a unified framework for integrating plant metabolic pathway knowledge across diverse data types and species, enabling new opportunities for hypothesis generation in plant natural product research across species from distributed biological knowledge. By combining curated metabolic pathway annotations with Semantic Web technologies, PlantMetWiki facilitates the exploration of biosynthetic pathways as interconnected, queryable graphs. This approach supports collaborative discovery by allowing researchers to identify gaps in pathway knowledge, compare biosynthetic routes across taxa, transfer knowledge from well-characterized species to less-studied taxa, and link chemical occurrence data to underlying biosynthetic mechanisms. The resulting graph captures both shared and species-specific aspects of metabolism, providing a reusable framework for studying pathway evolution, annotation transfer, and specialized metabolism across the plant kingdom. In addition, the availability of a SPARQL endpoint allows seamless integration with visualization and analysis platforms such as Cytoscape ^19^, facilitating the exploration of metabolic and regulatory networks in combination with user-provided omics data.

These findings demonstrate the power of interoperability for generating biologically meaningful insights: by making pathway, taxonomic, genomic, and chemical information queryable within a common framework, PlantMetWiki enables systematic assessment of annotation completeness across complementary resources. Federated queries allow researchers to move seamlessly from biological activity and toxicology to biosynthetic origin, genomic context, and species distribution, and address completeness and curation of knowledge gaps.

### Limitations: annotation quality of source databases, genome versioning, identifier mapping, and external endpoint availability

Several limitations should be considered when using and interpreting the current resource. First, PlantMetWiki inherits limitations from its source databases. Pathway completeness, species coverage, and identifier consistency ultimately depend on the underlying curation available in PlantCyc, MIBiG, plantiSMASH, and external reference resources. Second, cross-species inference to resolve annotation gaps is limited to truly shared reaction steps across species, which is known to be only partial due to the intrinsic fast-evolving nature of plant specialized metabolism ^37,38^. Third, integration across genomic resources remains constrained by differences in genome annotation systems and identifier schemes: incompatible annotation frameworks can limit automated cross-resource mapping ^39^. For instance, for tomato (*Solanum lycopersicum*), direct linking of PlantCyc data with plantiSMASH and MIBiG annotations was not implemented because plantiSMASH uses the NCBI RefSeq genome annotation (GCF_036512215.1, generating LOC identifiers and gene symbols; BioProject PRJDB16644), while PlantCyc uses the ITAG annotation ^40^ (generating Solyc identifiers hosted in SolGenomics Network ^41^). These two annotation systems define different gene models for the same genome sequence and cannot be directly matched. BridgeDb can support mapping annotation across resources, but the capacity of such tools needs to be constantly updated to represent different sources and taxa, and it cannot bridge across truly different annotations, such as gene annotations coming from different reference genomes, which is known as one of the biggest challenges limiting the transferability of findings and integration of omics experiments in the plant sciences field ^42^. Maintaining and extending current mapping tools to additional species and genome annotations will be important for improving interoperability across plant genomic resources. Additionally, the dependence on genome annotations introduces challenges related to versioning and consistency, particularly in plant systems where genome assemblies and annotations are frequently updated. Addressing this issue requires adherence to FAIR data principles and explicit tracking of genome annotation versions, for example, through standardized references to GenBank assemblies, to ensure reproducibility and transparency. PlantMetWiki knowledge graph provides a flexible framework to manage these evolving datasets but does not eliminate the need for careful data curation.

Federated analyses depend on the continued availability and interoperability of external resources. While LOD approaches reduce information duplication, they also rely on external services to maintain stable identifiers and accessible query interfaces. This limitation is particularly apparent for plantiSMASH and MIBiG, whose biological content is currently accessible primarily through web interfaces rather than semantically modelled LOD, limiting the discoverability of the information contained in these resources. As a result, PlantMetWiki can link users to biosynthetic gene clusters, but cannot yet perform deeper federated queries across cluster-associated genes, metabolites, taxonomic annotations, or chemical structures. The utility of federated analyses remains dependent on the continued availability of public SPARQL endpoints such as Wikidata and the NCBI Taxonomy project ^43^. These limitations underscore the importance of continued adoption of FAIR principles, identifier harmonization, and LOD technologies across plant biology resources. Expanding the ecosystem of queryable resources, for example, through public mirrors of databases such as ChEMBL or the conversion of domain-specific datasets into LOD formats, will further enhance the utility of Semantic Web platforms like PlantMetWiki.

### Conclusions and Outlook: PlantMetWiki as a FAIR infrastructure for pathway knowledge

PlantMetWiki provides a foundation for a continuously evolving, community-driven knowledge base for plant biosynthesis. The integration of additional biological and omics layers will further enhance the ability to model complex biosynthetic systems and generate testable hypotheses. The PlantMetWiki framework can be extended to incorporate additional layers of biosynthetic regulation, including transcription factor networks and other regulatory annotations, developmental stages, phenotypes, gene homology, or orthologous relationships, as well as alternative genome mining or pathway-finding tools beyond plantiSMASH. Experimental components such as tissues and developmental stages can be integrated to advance the interpretation of omics and multi-omics layers at increased resolution, as it has been demonstrated across paired omics studies ^29,44,45^. Such integration highlights the potential of PlantMetWiki as a central hub for linking pathway knowledge with emerging computational tools for biosynthetic discovery relying on multi-omics, such as NPLinker ^46^ or MEANtools ^47^. Furthermore, PlantMetWiki has the potential to connect to other knowledge graphs in the plant sciences, such as KnetMiner ^48^, for gene and gene networks, AgroLD ^49^, for molecular and phenotypic information, PlantConnectome ^50^, for plant science literature, and Stress Knowledge Map ^51^, for plant stress responses. Complementary to these resources, PlantMetWiki shows the potential of Semantic Web technologies and LOD to transform various types of plant data into actionable knowledge for hypothesis formulation and validation.

The modular architecture of PlantMetWiki enables iterative updates and continuous integration of new data sources, including future releases of the source databases, Plant Metabolic Network, BridgeDb identifier mappings, plantiSMASH predictions, and MIBiG annotations. Furthermore, the use of federated queries enhances interoperability by allowing dynamic connections to external resources that are regularly updated, such as Wikidata, ChEBI, PubMed, PubChem, and others. This approach can help address common challenges in data integration, such as incomplete cross-referencing of literature identifiers or missing metabolite annotations, by linking distributed knowledge across databases without requiring centralized storage. Future work should focus on expanding species coverage beyond current PlantCyc annotations and enabling joint querying of experimentally validated and computationally predicted biosynthetic pathways.

The entire workflow underlying PlantMetWiki is transferable to other pathway collections from the Pathway Tools ecosystem, such as MetaCyc, BioCyc organism databases: the conversion pipeline operates on standard BioCyc flat-file exports and produces FAIR RDF representations with machine-readable metadata and validation procedures. PlantMetWiki therefore represents both a biological resource and a reusable methodology for transforming pathway databases into a fully deployed RDF resource and interoperable knowledge graphs.

PlantMetWiki demonstrates how plant metabolic pathway knowledge can be represented as a FAIR multispecies knowledge graph that supports integration across pathway, taxonomic, genomic, and chemical domains. By combining semantic representation with interoperable identifiers and federated querying, the resource enables comparative analyses, systematic assessment of annotation completeness, and integration with complementary biological knowledge sources. Beyond the biological content itself, PlantMetWiki provides a reproducible and transferable workflow for transforming pathway databases into FAIR knowledge graphs, supporting broader reuse of pathway knowledge within the life sciences LOD ecosystem

## Materials and Methods

### Generation of cross-species plant pathway representations

Pathway and biochemical information from PlantCyc 17.0.0 (Plant Metabolic Network; December 2025 release) is retrieved in BioCyc flat-file format and converted into GPML2021 pathway representations using a custom open-source data extraction workflow (see Data availability section). Data extraction proceeds bottom-up, starting from each pathway record. For each pathway, constituent reactions, metabolites, proteins, genes, enzyme complexes, regulatory interactions, and literature references are retrieved and represented in GPML with the custom Cyc2Wiki pipeline (https://github.com/pathway-lod/Cyc_to_wiki). Reactions present in PlantCyc but not assigned to a pathway were exported as standalone reaction representations to preserve the coverage of all biochemical transformations. The resulting GPML files constitute the foundation of the PlantMetWiki knowledge graph and provide a reproducible intermediate format for downstream semantic conversion. The reaction layout was computed using a grid-based algorithm. A primary compound was identified for each reaction to serve as the spatial anchor, and positions for all DataNode elements were calculated relative to it, arranged in three layers: genes at the top, proteins and enzyme complexes in the middle, and metabolites (substrates/products) at the bottom.

Taxonomy annotation was applied at two levels. At the pathway level, all 2,478 GPML files carry organism="Viridiplantae" (NCBI Taxonomy ID: 33090). At the entity level, individual GeneProduct and Protein DataNodes carry <Annotation type="Taxonomy"> elements, linking each entity to its source organism via NCBI Taxonomy. Because the gene flat file in PlantCyc does not contain a SPECIES field, gene taxon annotations were propagated from the linked protein product via the PRODUCT field. Species-level annotations on protein complex DataNodes (CPLX-type entries) are handled separately. Enzyme complexes are rendered as <Group type="Complex"> elements in GPML rather than individual DataNodes. Because the GPML2021 XSD does not permit <ANNOTATIONREF> elements on <GROUP> elements, the species annotation of a complex record cannot be attached directly to the Group element. Instead, the SPECIES field of the complex record is propagated to its component monomer DataNodes during the build: if a monomer carries no independent SPECIES annotation in proteins.dat, it inherits the parent complex’s annotation. However, component monomers only appear as standalone DataNodes in GPML when they directly catalyze at least one reaction: monomers that serve exclusively as structural subunits are never emitted as independent DataNodes.

Quality assurance is performed before each build with a custom validation step that checks the consistency of protein–gene relationships in the PlantCyc flat files. The primary check flags genes whose protein products are annotated to different NCBI taxa and spots potential curation issues in the source database. The validator reports these issues and skips these entries, preventing incorrect annotations from propagating silently into the GPML output (Supplementary Table S3).

Citations from PlantCyc (pubs.dat source flat file) are transferred to the GPML as <CITATION> elements following the GPML2021 model. Each citation carries an Xref with a PubMed ID, DOI, or PlantCyc-internal identifier as fallback, in order of priority. <CITATIONREF> elements on individual DataNodes and Interactions link each entity to its supporting literature. Identifier provenance is ensured by preserving the original PlantCyc UNIQUE-ID values in <Property key="UniqueID"> elements within each DataNode, maintaining a direct traceable link to the source PMN records. GPML schema validation involves all generated GPML files being validated against the GPML2021 XML Schema Definition using an automated GitHub Actions workflow at every commit (https://github.com/PathVisio/GPML/blob/master/GPML2021/GPML2021.xsd).

### RDF conversion, semantic enrichment, and BridgeDB mapping

RDF conversion of the GPML files was performed using the gpml2rdf Java tool (v4.0.4, org.wikipathways.wp2rdf.CreateRDF), which implements the WikiPathways RDF model and uses libGPML v4.0.4 ^52^ for GPML parsing. The full pipeline is available at https://github.com/pathway-lod/gpml-to-rdf. Before conversion, each GPML file is assigned a stable numeric identifier (PC1–PC1162 for pathways; RC1–RC1316 for reactions) using Groovy renaming scripts, decoupling the RDF identifiers from the PlantCyc pathway IDs while preserving the original identifiers as properties inside the graph.

Core RDF conversion is parallelized across GPML files using GNU Make (make -j), with each file converted independently. The converter produces two representations per file: a WikiPathways RDF (WPRDF) file following the “wp:Pathway” semantic model, and a GPML RDF (GPMLRDF) file capturing the raw GPML structure, including all the PlantCyc-derived information. All Internationalized Resource Identifiers (IRIs) use the base domain rdf-plantmetwiki.bioinformatics.nl, generating pathway IRIs of the form “http://rdf-plantmetwiki.bioinformatics.nl/pathways/PC{N}_r{version}”. BridgeDB (build 20260102; derived from HMDB, ChEBI, and Wikidata) metabolite identifier mapping adds independent information via “wp:bdb*” predicates (wp:bdbChEBI, wp:bdbHmdb, wp:bdbWikidata, wp:bdbPubChem, wp:bdbInChIKey, wp:bdbKeggCompound, wp:bdbLipidMaps, wp:bdbChemspider). The PlantCyc-sourced dc:identifier/InChIKey stays untouched, resulting in an augmentation with preserved provenance to distinguish PlantCyc-curated annotations from BridgeDB-derived ones.

Downstream enrichment was performed sequentially in Python via two plugin scripts that extend the same IRI subjects as the core graph. First, taxonomy information is mapped to the pathway model by (1) taking the <Annotation type="Taxonomy"> and <ANNOTATIONREF> elements from each GPML file and attaching “wp:organism” Viridiplantae (NCBI:33090) to the pathway IRI, and (2) attaching it to each GeneProduct, Protein, and Metabolite DataNode IRI with their species-specific NCBI taxon resulting from PlantCyc. A “foaf:page” triple is also added to the main pathway IRI, linking it to the corresponding clickable PlantCyc entry in the Plant Metabolic Network browser (https://pmn.plantcyc.org/pathway?orgid=PLANT&id={PlantCycID}). Second, the additional properties from the PlantCyc GPMLs that are not semantically modelled in the core PlantMetWiki data model (interoperable with WikiPathways) are preserved as a separate “graph/gpml-properties-extra” layer representing PlantCyc-specific <Property key="" value=""> elements from Pathway, DataNode, and Interaction elements as “pmw:gpmlProperty” blank nodes. For the UniqueID property, this graph also contains “pmw:plantcycId” (the original PlantCyc identifier string), “dcterms:source” (linking to the stable https://identifiers.org/plantcyc/{id}), and clickable browser links to PMN via FOAF RDF Vocabulary ^53^ “foaf:page” (Supplementary Table S4).

RDF validation is performed at three levels using RDFlib (https://github.com/RDFLib/rdflib/): (1) syntax parsing of every individual TTL file output by the converter and plugins; (2) content checks on the aggregated taxonomy and properties bundles for required predicates and minimum triple counts; (3) pattern scanning of the large core graph bundle and full parsing of the VoID file, verifying that at least four “void:Dataset” instances are present. The complete RDF dataset is uploaded to the PlantMetWiki community on Zenodo via the Zenodo API, with metadata and license information derived from the VoID header. The resulting RDF is structured as four named graphs: (1) core pathways, represent the pathway model for all 1,162 PlantCyc pathways (2) core reactions, contain the model for all 1,316 individual PlantCyc-derived reactions (3) taxonomy extra, contains taxonomic annotations (wp:organism) to gene, protein, and compound nodes, including “foaf:page” links. (4) properties extra, contains links to PlantCyc identifiers, stable pathway Uniform Resource Identifiers (URIs), and key-value properties.

### Extending pathway knowledge with crosslinks to biosynthetic gene clusters

The PlantCyc-derived pathway RDF is extended with crosslinks to two BGC databases: MIBiG (curated BGCs, ^22^) and plantiSMASH 2.0 (predicted plant BGCs, ^26^), with a custom open-source pipeline (see Data availability). BGC gene cluster membership data from both sources was converted to RDF Turtle format following a BGC-first data model. Each cluster is represented as a “pmw:BiosyntheticGeneCluster” resource with its constituent genes linked via the OBO Relations Ontology predicate RO:0000051 (has_part). The “dcterms:source” predicates record whether each cluster originates from MIBiG or plantiSMASH. Gene identifiers are normalized to identifiers.org/tair.name/ IRIs, consistent with the WikiPathways/PlantMetWiki pathway RDF.

For *A. thaliana*, BGC-to-pathway links are established by a direct SPARQL join on the shared TAIR gene identifier: both the PlantCyc pathway graph and the plantiSMASH BGC graph use identifiers.org/tair.name/ IRIs for *A. thaliana* genes, enabling unambiguous matching without external identifier mapping services. From MIBiG, known *A. thaliana* BGCs and their genes are tagged to “wp:organismName” "*Arabidopsis thaliana*" and “wp:organism” ncbi:3702, representing tair.name IRIs that can connect to PlantCyc pathways. The other plant MIBiG BGCs that do not match *A. thaliana* are kept as “pmw:MemberIdentifier nodes” by design, as their identifiers (XP_/NP_ accessions etc.) can’t reliably be annotated to a species. Validation via RDFlib (https://github.com/RDFLib/rdflib/) of the final Turtle files is implemented. The RDF datasets are described using a VoID metadata file that records dataset provenance, triple counts, and licensing information (see Data availability section).

### Deployment, validation and accessibility

PlantMetWiki is available via a SNORQL user interface with example queries that can be customized; a tutorial and documentation are available. The pathway data and connected annotations are made available via a SPARQL endpoint which can be accessed by the PlantMetWiki SPARQL Explorer, a SNORQL User Interface (UI) including a panel with example queries. SNORQL is a query editor that offers syntax highlighting for writing and executing SPARQL queries directly on our existing SPARQL endpoint, mirroring the usage of WikiPathways ^54^. The example queries auto-populate the SPARQL Explorer and can be adapted by users directly on the webserver, or in a customizable GitHub repository (https://github.com/pathway-lod/SPARQLQueries). This repository stores queries in folders allowing new users to navigate through our example queries panel with more ease and to generate permalinks to their queries for reproducibility of the results. This setup allows to collaborate on the creation of queries over topics, communities, collaborations functionality, or external data sources for federated queries. Links of PlantCyc pathways with BGCs can be browsed with the “BGC Crosslinks” menu.

The example queries provided in the UI include federated queries using the QLever ^55^ search engine for quicker and more reliable access to Wikidata (https://qlever.cs.uni-freiburg.de/api/wikidata). A tutorial is included to guide users on the creation and customization of SPARQL queries (see Data availability section). These can be customized and the user can point the UI to other repositories containing custom queries.

To enable local taxon label resolution and taxonomic reasoning without a runtime dependency on an external federation endpoint, the NCBITaxon ontology (release v2026-05-13, OBO Foundry, CC0 1.0 Universal) ^43^ is loaded from http://purl.obolibrary.org/obo/ncbitaxon.owl into a dedicated “graph/ncbitaxon” named graph. The pathway IRIs already use the Open Bio-Ontologies (OBO) Foundry ^56^ IRI scheme (http://purl.obolibrary.org/obo/NCBITaxon_<ID>), so taxon labels are resolved through a simple cross-graph SPARQL join without any IRI translation. ROBOT ^30^ (method MIREOT) is used to downsize the NCBITaxon ontology and extract all PlantCyc-derived taxa plus their ancestors up the lineage to the ontology root. This provides the “rdfs:label” of every taxon in the lineage and the complete subClassOf hierarchy from leaf to root, needed for tree navigation. The taxonomy-extra dataset’s VoID metadata explicitly declares the OBO Foundry NCBITaxon namespace as a “void:vocabulary” and links to the full ontology release via “dcterms:references”, making the provenance of all taxon IRIs machine-readable. The MIREOT extraction method and ROBOT tooling are documented in the dataset description, enabling reproducibility of the local SPARQL endpoint setup.

The VoID metadata files describe the four named graphs in the Virtuoso triple store as “void:Dataset” instances, and include record dataset titles, versioning, triple counts, publisher, licensing and provenance (Zenodo DOI, build timestamp, PlantCyc version) metadata. Every “void:Dataset” (core pathway RDF, taxonomy-extra, properties-extra, BGC mibig-4.0/plantismash-v2, and NCBITaxon ROBOT subset) carries “dcterms:publisher”, pointing to a “foaf:Organization”, plus “dcterms:modified/“pav:createdOn”” to record freshness. VoID discoverability: the combined VoID description is served at the “well-known VoID” (RFC 8615 URI scheme) in addition to being queryable as a named graph via SPARQL, so harvesters can locate dataset metadata without prior knowledge of the endpoint’s graph structure. Each dataset includes a “foaf:page” link to documentation of the underlying “wp:/pmw”: RDF class structure, clarifying how pathway, taxonomy, and gene-cluster resources relate to the data model. Biosynthetic gene cluster resources (pmw:BiosyntheticGeneCluster) include “rdfs:seeAlso” links to their source records in plantiSMASH and MIBiG, and a “void:Linkset” connects BGC genes to PlantMetWiki pathway genes. Licensing is represented via “dcterms:license/“dcterms:rights”” to specify reuse terms per dataset (PMN Open Database License for pathway data, CC-BY-4.0 for MIBiG, GPL-3.0 for plantiSMASH). All VoID dataset metadata is loaded into a named graph, which automatically exposes it alongside the data graphs in Virtuoso.

## Data availability

All relevant data are deposited and versioned in Zenodo at https://zenodo.org/communities/plantmetwiki/, including relevant license and copyright information. Tutorial pages are available at https://pathway-lod.github.io/SPARQLTutorials/. Jupyter notebooks containing the steps to reproducibly generate the figures are available on GitHub at https://github.com/pathway-lod/gpml-to-rdf/tree/main/notebooks. PlantMetWiki is listed on bio.tools at https://bio.tools/plantmetwiki. Further project metadata is available on the Nanodash platform (Knowledgepixels) at https://w3id.org/spaces/plantmetwiki.

## Code availability

All relevant code to this manuscript is deposited in GitHub at https://github.com/pathway-lod and deposited in versioned releases in Zenodo at https://zenodo.org/communities/plantmetwiki/

## Supplementary tables

**Supplementary Table S1.** Summary of taxa per NCBI ID represented in the GPML files.

**Supplementary Table S2.** Summary of taxonomic information captured by PlantMetWiki in the GPML representation.

**Supplementary Table S3.** Validation pipeline of GPML to RDF conversion.

**Supplementary Table S4.** Summary of extra properties captured by the third graph and not semantically modelled.

**Supplementary Table S5.** Summary of BGC mapping to PlantMetWiki from MIBiG and plantiSMASH.

**Supplementary Table S6**. Overview of the PlantMetWiki Virtuoso triplestore.

**Supplementary Table S7**. PlantMetWiki metabolites absent from Wikidata.

## Author contributions

Elena Del Pup: Conceptualization, Data curation, Formal analysis, Funding acquisition, Investigation, Software, Writing – original draft. Max Muller: Data curation, Formal analysis, Software, Validation. Egon L. Willighagen: Conceptualization, Formal analysis, Methodology, Software, Writing – review & editing. Marvin Martens: Methodology, Software, Validation. Marnix H. Medema: Funding acquisition, Project administration, Supervision, Writing – review & editing. Denise Slenter: Conceptualization, Data curation, Formal analysis, Methodology, Software, Supervision, Validation, Writing – original draft, Writing – review & editing. Justin J.J. van der Hooft: Conceptualization, Funding acquisition, Project administration, Supervision, Writing – review & editing.

## Acknowledgements

We thank Sue Rhee and the Plant Metabolic Network curators for granting an open license that allows reuse of their data, and for their valuable feedback. We thank Simon Shaw and Mitja Zdouc for valuable advice on linking biosynthetic gene cluster information to PlantMetWiki. We also thank Arjan Draisma and Harm Nijveen for advice on hosting the database and website for the SPARQL Explorer. We thank Adriano Rutz for input on the use of WikiData and LOTUS, and Leen Abraham for advice on integrating plant reference genomes. Finally, we thank the Open Science and Open Data community for building and maintaining the resources that PlantMetWiki can connect to for its queries.

## AI usage statement

During the preparation of this work, the authors used ChatGPT for brainstorming and obtaining feedback on the manuscript; Claude and Claude Code for coding assistance and debugging; GitHub Copilot to aid in code review; and Grammarly for copyediting. After using these tools, the authors reviewed and edited the content as needed and take full responsibility for the content.

## References

1. Huang, X.-Q. & Dudareva, N. Plant specialized metabolism. Current Biology 33, R473–R478 (2023).

2. Domingo-Fernández, D. et al. Modern drug discovery using ethnobotany: A large-scale cross-cultural analysis of traditional medicine reveals common therapeutic uses. iScience 26, 107729 (2023).

3. Owen, C., Steuernagel, B., Hall, N., Paige, B. & Osbourn, A. Mind the gap: the challenges and opportunities for genomics-driven harnessing of plant metabolic diversity for therapeutic applications. Current Opinion in Biotechnology 99, 103510 (2026).

4. Greco, M., Caminada, G., Coculo, D. & Lionetti, V. From Waste to Defense: Agro-Industrial Byproducts as Sources of Biopesticides and Bioelicitors for Crop Protection. J. Agric. Food Chem. 74, 10624–10644 (2026).

5. Ji, W., Osbourn, A. & Liu, Z. Understanding metabolic diversification in plants: branchpoints in the evolution of specialized metabolism. Philosophical Transactions of the Royal Society B: Biological Sciences 379, 20230359 (2024).

6. Rutz, A. et al. The LOTUS initiative for open knowledge management in natural products research. eLife 11, e70780 (2022).

7. Sorokina, M., Merseburger, P., Rajan, K., Yirik, M. A. & Steinbeck, C. COCONUT online: Collection of Open Natural Products database. Journal of Cheminformatics 13, 2 (2021).

8. Mildau, K. et al. Effective data visualization strategies in untargeted metabolomics. Natural Product Reports 10.1039/D4NP00039K (2025) doi:10.1039/D4NP00039K.

9. Saurabh Singh, K., Hooft, J. J. J. van der, Wees, S. C. M. van & H. Medema, M. Integrative omics approaches for biosynthetic pathway discovery in plants. Natural Product Reports 39, 1876–1896 (2022).

10. Uauy, C. et al. Challenges of translating Arabidopsis insights into crops. Plant Cell 37, koaf059 (2025).

11. Scossa, F., Bulut, M., Naake, T., D’Auria, J. C. & Fernie, A. R. Convergence and parallelism in the evolution of plant metabolism. Journal of Integrative Plant Biology 68, 1013–1031 (2026).

12. Schoch, C. L. et al. NCBI Taxonomy: a comprehensive update on curation, resources and tools. Database (Oxford) 2020, baaa062 (2020).

13. Hawkins, C. et al. Plant Metabolic Network 16: expansion of underrepresented plant groups and experimentally supported enzyme data. Nucleic Acids Res 53, D1606–D1613 (2025).

14. Demir, E. et al. The BioPAX community standard for pathway data sharing. Nat Biotechnol 28, 935–942 (2010).

15. Karp, P. D. et al. Pathway Tools version 23.0 update: software for pathway/genome informatics and systems biology. Brief Bioinform 22, 109–126 (2021).

16. Kutmon, M. et al. PathVisio 3: An Extendable Pathway Analysis Toolbox. PLOS Computational Biology 11, e1004085 (2015).

17. Waagmeester, A. et al. Using the Semantic Web for Rapid Integration of WikiPathways with Other Biological Online Data Resources. PLOS Computational Biology 12, e1004989 (2016).

18. Agrawal, A. et al. WikiPathways 2024: next generation pathway database. Nucleic Acids Research 52, D679–D689 (2024).

19. Shannon, P. et al. Cytoscape: A Software Environment for Integrated Models of Biomolecular Interaction Networks. Genome Res. 13, 2498–2504 (2003).

20. Waagmeester, A. et al. Wikidata as a knowledge graph for the life sciences. Elife 9, e52614 (2020).

21. Hanumappa, M. et al. WikiPathways for plants: a community pathway curation portal and a case study in rice and arabidopsis seed development networks. Rice 6, 14 (2013).

22. Zdouc, M. M. et al. MIBiG 4.0: advancing biosynthetic gene cluster curation through global collaboration. Nucleic Acids Research 53, D678–D690 (2025).

23. Itkin, M. et al. Biosynthesis of Antinutritional Alkaloids in Solanaceous Crops Is Mediated by Clustered Genes. Science 341, 175–179 (2013).

24. Franke, J. et al. Gene Discovery in Gelsemium Highlights Conserved Gene Clusters in Monoterpene Indole Alkaloid Biosynthesis. ChemBioChem 20, 83–87 (2019).

25. Li, Y. et al. Subtelomeric assembly of a multi-gene pathway for antimicrobial defense compounds in cereals. Nat Commun 12, 2563 (2021).

26. Del Pup, E. et al. plantiSMASH 2.0: Improvements to Detection, Annotation, and Prioritization of Plant Biosynthetic Gene Clusters. Journal of Molecular Biology 169798 (2026) doi:10.1016/j.jmb.2026.169798.

27. Slenter, D. N. et al. WikiPathways: a multifaceted pathway database bridging metabolomics to other omics research. Nucleic Acids Res 46, D661–D667 (2018).

28. van Iersel, M. P. et al. The BridgeDb framework: standardized access to gene, protein and metabolite identifier mapping services. BMC Bioinformatics 11, 5 (2010).

29. Wolters, F. C. et al. Pairing omics to decode the diversity of plant specialized metabolism. Current Opinion in Plant Biology 82, 102657 (2024).

30. Jackson, R. C. et al. ROBOT: A Tool for Automating Ontology Workflows. BMC Bioinformatics 20, (2019).

31. Owen, C. plantiSMASH-2.0.4 database (with internal cyclopeptide repeat detection only). 10.5281/zenodo.1747281 (2025) doi:10.5281/zenodo.17472815.

32. Alexander, K., Cyganiak, R., Hausenblas, M. & Zhao, J. Describing linked datasets with the VoID vocabulary. in (2011).

33. Leveau, A. et al. Towards take-all control: a C-21β oxidase required for acylation of triterpene defence compounds in oat. New Phytologist 221, 1544–1555 (2019).

34. Malik, A. et al. ChEBI: re-engineered for a sustainable future. Nucleic Acids Res 54, D1768–D1778 (2026).

35. Huang, R., O’Donnell, A. J., Barboline, J. J. & Barkman, T. J. Convergent evolution of caffeine in plants by co-option of exapted ancestral enzymes. Proc Natl Acad Sci U S A 113, 10613–10618 (2016).

36. Çiçek, S. S., Mangoni, A., Hanschen, F. S., Agerbirk, N. & Zidorn, C. Essentials in the acquisition, interpretation, and reporting of plant metabolite profiles. Phytochemistry 220, 114004 (2024).

37. Weng, J.-K., Philippe, R. N. & Noel, J. P. The rise of chemodiversity in plants. Science 336, 1667–1670 (2012).

38. Dadras, A. et al. Accessible versatility underpins the deep evolution of plant specialized metabolism. Phytochem Rev 24, 13–26 (2025).

39. Monnahan, P. J. et al. Using multiple reference genomes to identify and resolve annotation inconsistencies. BMC Genomics 21, 281 (2020).

40. Sato, S. et al. The tomato genome sequence provides insights into fleshy fruit evolution. Nature 485, 635–641 (2012).

41. Fernandez-Pozo, N. et al. The Sol Genomics Network (SGN)—from genotype to phenotype to breeding. Nucleic Acids Res 43, D1036–D1041 (2015).

42. Roeder, A. H. K. et al. Lost in translation: What we have learned from attributes that do not translate from Arabidopsis to other plants. Plant Cell 37, koaf036 (2025).

43. Federhen, S. The NCBI Taxonomy database. Nucleic Acids Res 40, D136–D143 (2012).

44. Barrios, W. E. B. et al. Integrated transcriptome–metabolome analyses reveal regulatory networks underlying soluble solids accumulation in Capsicum chinense fruits. Plant J 127, e71020 (2026).

45. Jeon, J. E. et al. A Pathogen-Responsive Gene Cluster for Highly Modified Fatty Acids in Tomato. Cell 180, 176–187.e19 (2020).

46. Geng, C. et al. NPLinker 2: a modular and customizable framework for paired omics analyses. Journal of Open Source Software 11, 9528 (2026).

47. Singh, K. S. et al. MEANtools integrates multi-omics data to identify metabolites and predict biosynthetic pathways. PLOS Biology 23, e3003307 (2025).

48. Hassani-Pak, K. et al. KnetMiner: a comprehensive approach for supporting evidence-based gene discovery and complex trait analysis across species. Plant Biotechnology Journal 19, 1670–1678 (2021).

49. Larmande, P. et al. AgroLD: a knowledge graph for the plant sciences. BMC Genom Data 26, 73 (2025).

50. Lim, S. C. et al. PlantConnectome: A knowledge graph database encompassing >71,000 plant articles. Plant Cell 37, koaf169 (2025).

51. Bleker, C. et al. Stress Knowledge Map: A knowledge graph resource for systems biology analysis of plant stress responses. Plant Communications 5, 100920 (2024).

52. Pico, A., et al. libGPML. Zenodo 10.5281/ZENODO.18073687 (2025).

53. Brickley, D. & Miller, L. FOAF Vocabulary Specification. https://xmlns.com/foaf/spec/20140114.html.

54. Martens, M. et al. WikiPathways: connecting communities. Nucleic Acids Res 49, D613–D621 (2020).

55. QLever | Proceedings of the 2017 ACM on Conference on Information and Knowledge Management. https://dl.acm.org/doi/10.1145/3132847.3132921.

56. Smith, B. et al. The OBO Foundry: coordinated evolution of ontologies to support biomedical data integration. Nat Biotechnol 25, 1251 (2007).

