## Supplementary Figures for "PlantMetWiki: a FAIR knowledge graph for plant metabolic pathway cross-species representation and integration"

[1] Bioinformatics Group, Wageningen University, Wageningen, the Netherlands. [2] Department of Translational Genomics, NUTRIM, Maastricht University, Maastricht, the Netherlands.[3] Institute of Biology, Leiden University, Leiden, the Netherlands. [4] Department of Advanced Computing Sciences, DACS, Maastricht University, Maastricht, the Netherlands. [5] Institute of Data Science, IDS, Maastricht University, the Netherlands [6]Department of Biochemistry, University of Johannesburg, Johannesburg 2006, South Africa.

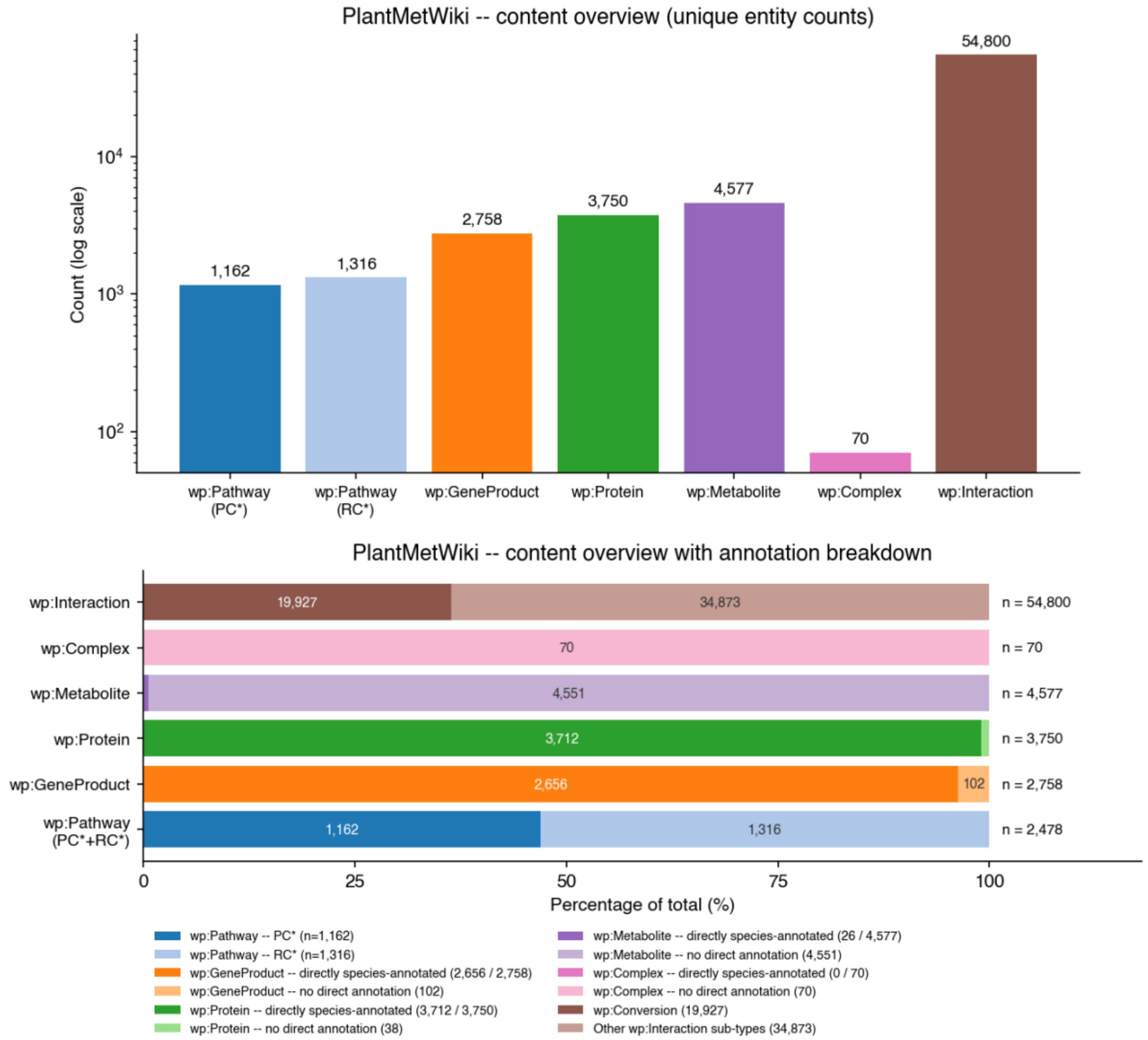


Figure S1. **Top panel: Overview bar chart.** Total unique counts of the main wp: content types in PlantMetWiki, on a logarithmic scale. Counts are true *distinct-entity* totals (COUNT(DISTINCT ?e) per class). **wp:Pathway (PC*)** and **wp:Pathway (RC*)**: pathways and reactions are both typed wp:Pathway by the gpml2rdf converter and are distinguished only by their IRI prefix. **wp:GeneProduct**, **wp:Protein**, **wp:Metabolite**, **wp:Complex**: DataNode entity types from the WikiPathways vocabulary (http://vocabularies.wikipathways.org/wp#). **wp:Interaction**: every instance of the parent interaction class (covers all typed edges: wp:Conversion, wp:Catalysis, wp:TranscriptionTranslation, wp:Inhibition, wp: Stimulation, wp:Binding, wp:ComplexBinding). **Bottom panel: PlantMetWiki core graph annotation coverage overview.** Proportional breakdown (100% per row) of each entity type by direct species-annotation status. Direct annotation: the entity carries a wp:organism triple in graph/gpml-taxonomy-extra, pointing to an NCBITaxon IRI (http://purl.obolibrary.org/obo/NCBITaxon_*). Dark colours: directly annotated; light colours: no direct wp:organism triple. **Genes** (wp:GeneProduct): 2,656 / 2,758 annotated (96 %). **Enzymes** (wp:Protein): 3,712 / 3,750 annotated (99 %). **Metabolites** (wp:Metabolite): only 26 / 4,577 carry a direct wp:organism; the rest are species-neutral in the source PlantCyc data. **Interactions**: split into wp:Conversion (biochemical reactions, brown) and all other wp: sub-types (light brown).

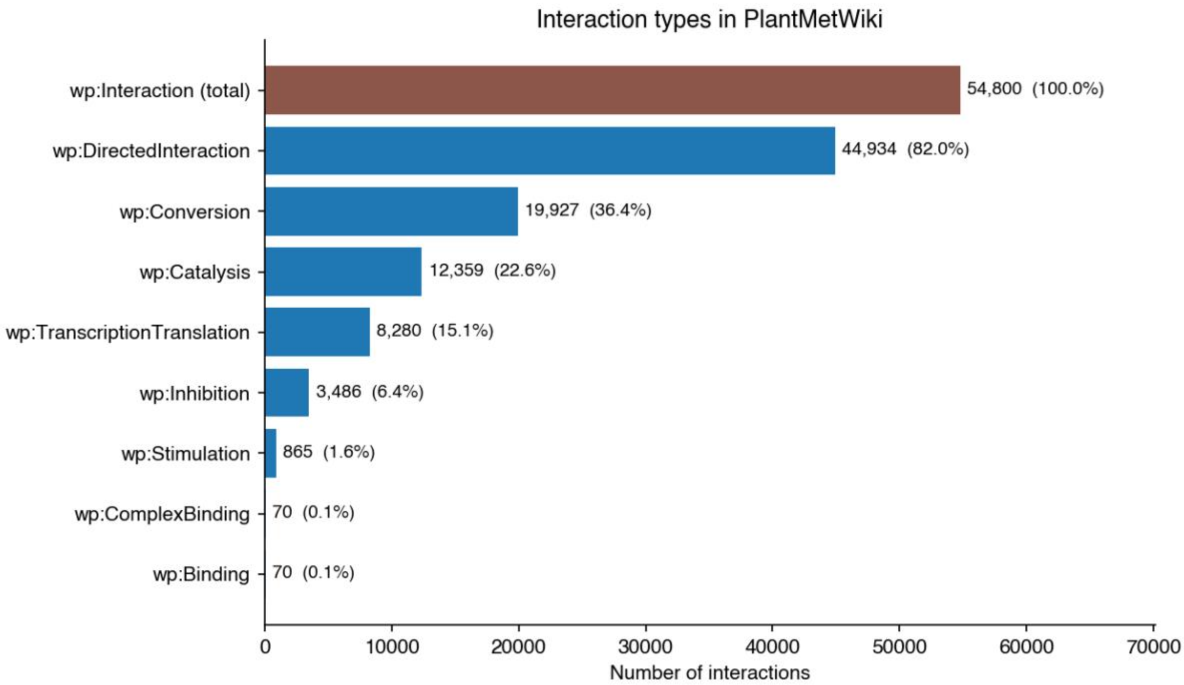


Figure S2. **Interaction type distribution.** wp:Interaction (parent class) instances shown alongside each specific wp: interaction sub-type (WikiPathways vocabulary, http://vocabularies.wikipathways.org/wp#) wp:Conversion: biochemical transformation connecting a metabolite substrate to a metabolite product. wp:Catalysis: link from an enzyme (wp:GeneProduct or wp:Protein) to the reaction it catalyses. wp:DirectedInteraction: generic directed edge (used for transport, regulatory flow, etc.). wp:TranscriptionTranslation: links gene, to transcript, to protein. wp:Inhibition, wp:Stimulation: negative or positive regulatory effects. wp:Binding or wp:ComplexBinding: physical interaction or complex formation. Sub-type counts sum to more than the parent count because edges can carry multiple wp: type labels.

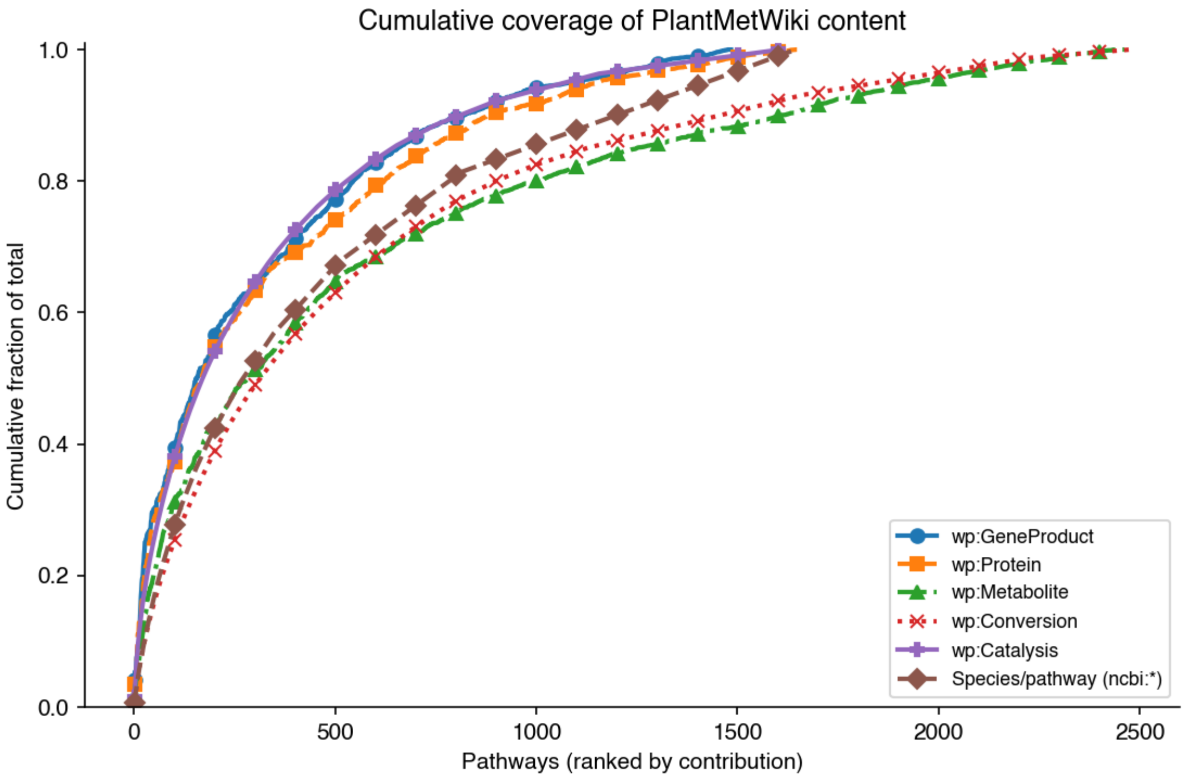


Figure S3. **PlantMetWiki core graph cumulative coverage curves.** For each metric, pathways are ranked by count (highest first) and the cumulative fraction of the dataset total is plotted against pathway rank. Pathway membership is encoded via dcterms:isPartOf; entity types are wp:GeneProduct, wp:Protein, wp:Metabolite, and wp:Conversion. wp:Conversion and wp:Catalysis interactions are minted with a pathway-specific IRI (each belongs to exactly one pathway), so their per-pathway counts already sum exactly to the unique totals (19,927 and 12,359 respectively). **Species/pathway** is calculated differently from the other five curves: for each pathway, COUNT(DISTINCT ?taxon) of species-specific wp:organism annotations on any wp:GeneProduct/wp:Protein node dcterms:isPartOf that pathway. A steep initial rise indicates concentration in a small number of large pathways. Genes and enzymes rise steeply because *Arabidopsis thaliana* (NCBITaxon_3702) pathways dominate the top ranks. The species-per-pathway curve rises more slowly, reflecting that taxonomic diversity is distributed across many medium-sized pathways. Curves that don't reach 100% (metabolites reach ~63%, enzymes ~95%) indicate the remaining unique entities have no recorded dcterms:isPartOf membership to any wp:Pathway-typed resource in this ranking.

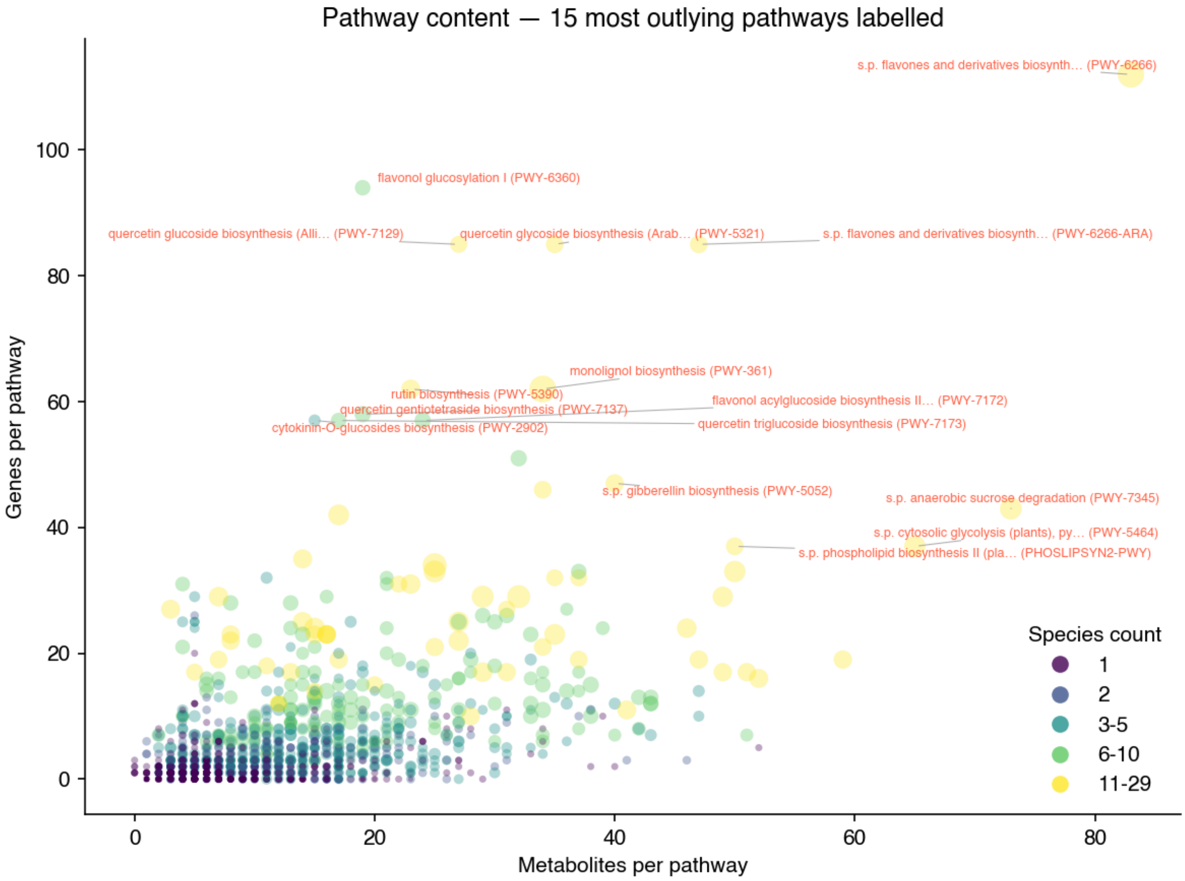


Figure S4. **PlantMetWiki core graph pathway content scatter plot with labelled outliers.** Same data and encoding as the scatter plot above. Red rings and labels mark the 15 wp:Pathway objects furthest from the point-cloud centroid (Euclidean distance in wp:Metabolite × wp:GeneProduct space). These outliers include large superpathways (aggregating many sub-pathways via dcterms:isPartOf) and compound-rich maps where wp:Metabolite nodes were imported without corresponding wp:GeneProduct annotations. Each label shows the pathway title followed by its PlantCyc identifier in parentheses (dcterms:identifier in graph/gpml-properties-extra, e.g. "PWY-6585", captured from the GPML <Property key="UniqueID"> during conversion: this is the code to look up the same pathway directly in PlantCyc.

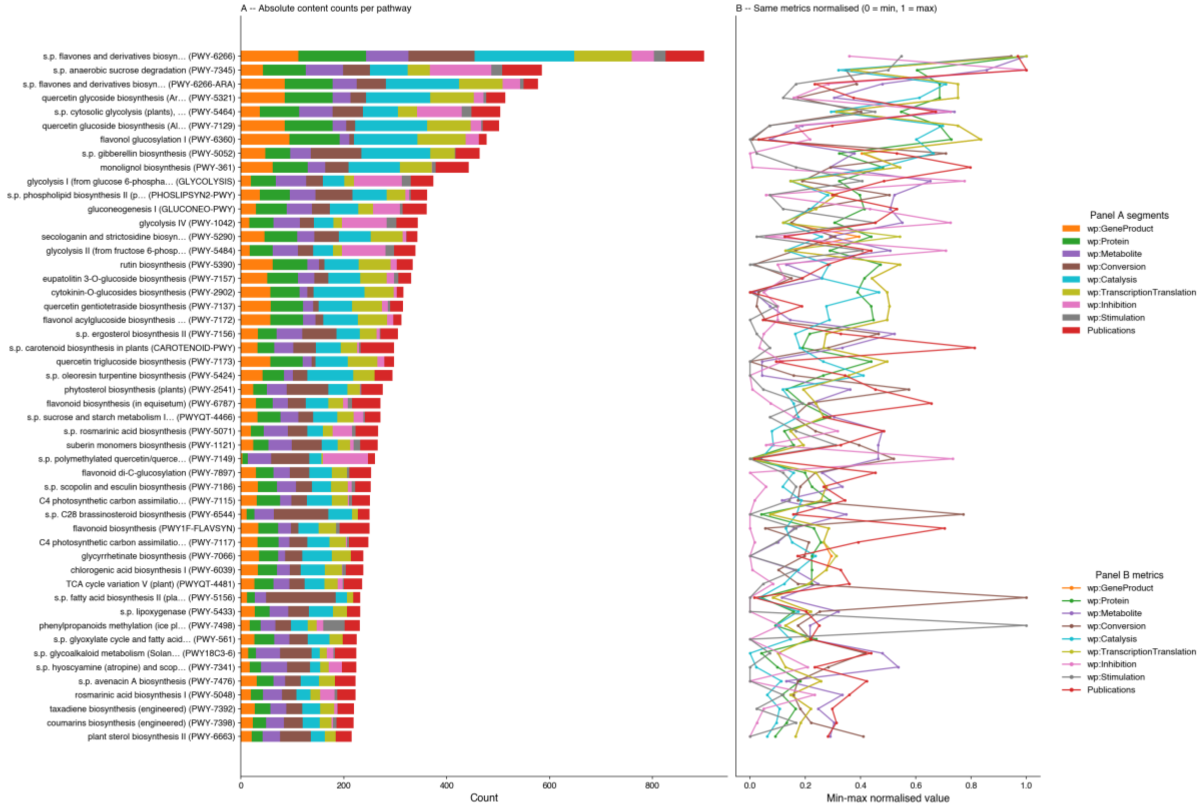


Figure S5. **Pathway content coverage (top 50 pathways).** Pathways ranked by total content (sum of all nine metrics below), most content-rich at top. PlantCyc PWY/RXN identifier shown in parentheses next to each pathway title (dcterms:identifier in graph/gpml-properties-extra). **(A)** Absolute counts, horizontal stacked
bars: wp:GeneProduct, wp:Protein, wp:Metabolite, wp:Conversion, wp:Catalysis, wp: TranscriptionTranslation, wp:Inhibition, wp:Stimulation (all dcterms:isPartOf that pathway), and linked publications (dcterms:references/cito:cites on the pathway itself). **(B)** Min-max normalization (0 = minimum, 1 = maximum across the top 50 pathways) of the same nine metrics. Unlike Figure 12 (species), there is no direct/indirect distinction here: every count is already scoped to a single pathway via direct RDF membership, so no catalysis-chain tracing is needed.

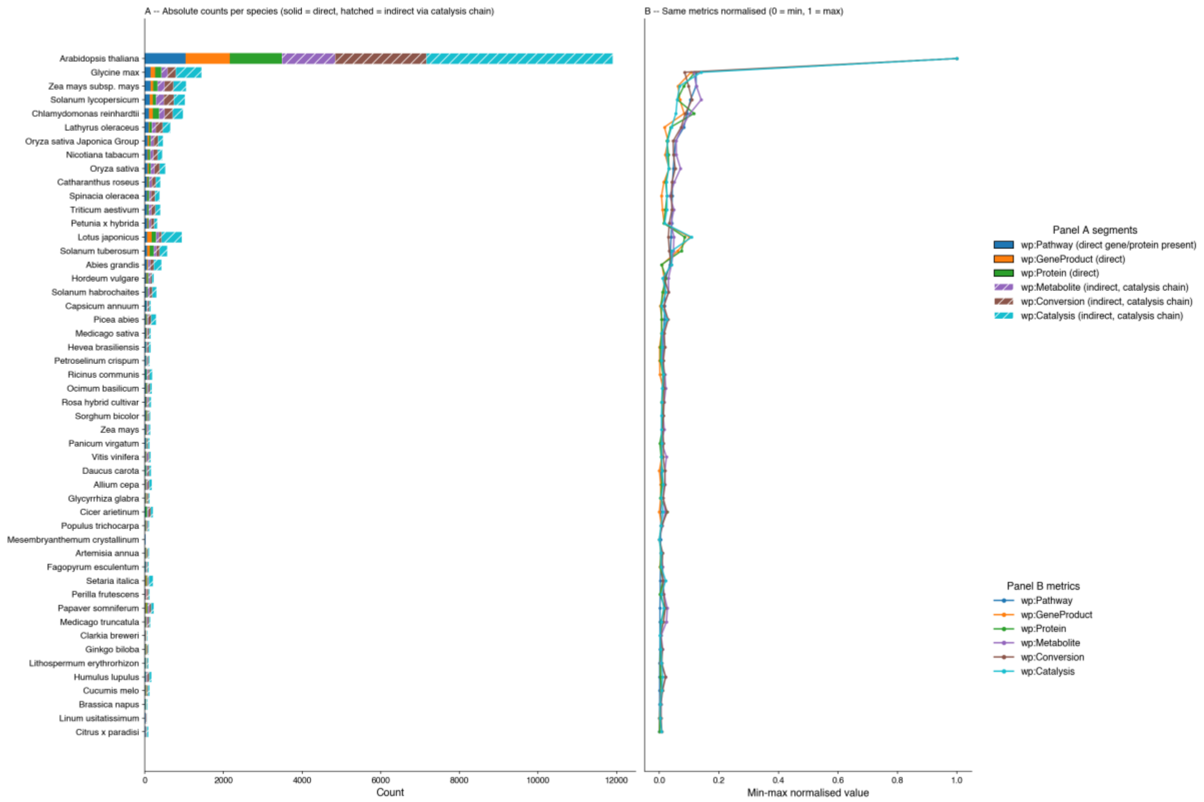


Figure S6. **PlantMetWiki core graph species coverage** (top 50 species). Both panels share the same vertical species axis (most pathway-rich at top), avoiding rotated category labels. **(A)** Absolute counts, horizontal stacked bars. Solid: wp:Pathway (containing a direct gene/protein of the species), wp:GeneProduct/wp:Protein with a direct wp:organism annotation. Hatched: wp:Metabolite, wp:Conversion, and wp:Catalysis attributed indirectly via the specific catalysis chain (wp:source/wp:target) the species' enzyme participates in (not broad pathway co-occurrence -- see Interpretation above). Publications are not included (pathway-level only, not chain-attributable). **(B)** Min-max normalisation (0 = minimum, 1 = maximum across the top 50 species) of: pathways, genes (wp:GeneProduct), enzymes (wp:Protein), metabolites (wp:Metabolite), conversions (wp:Conversion), and catalysis (wp:Catalysis). Normalisation reveals cross-species patterns independent of the large absolute differences driven by *Arabidopsis thaliana* (NCBITaxon_3702).

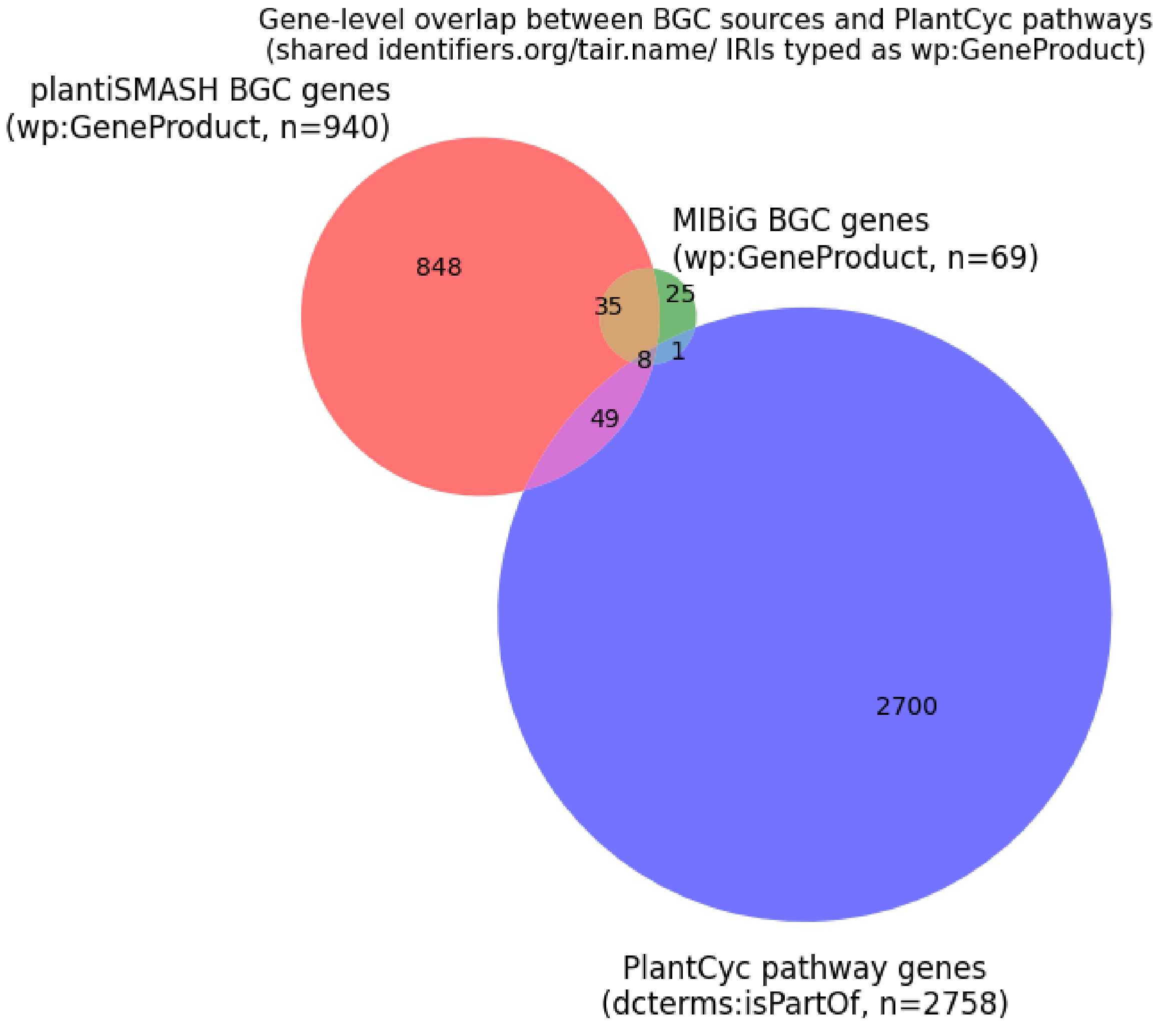


Figure S7. **Gene-level overlap between Biosynthetic Gene Cluster (BGC) sources and PlantCyc pathway annotations in PlantMetWiki.** Sets represent all wp:GeneProduct gene members identified by identifiers.org/tair.locus IRIs: plantiSMASH BGC genes (*n* = 940), MIBiG BGC genes (*n* = 69), and PlantCyc pathway genes (*n* = 2,758; all genes with dcterms:isPartOf a wp:Pathway in graph/pathways). Only wp:GeneProduct nodes (with identifiers.org IRIs) can form cross-graph joins; pmw:MemberIdentifier nodes (403 in MIBiG, 0 in plantiSMASH) are excluded. Of the 940 plantiSMASH BGC genes, 57 (6%) overlap with at least one PlantCyc pathway gene; for MIBiG, 9 of 69 (13%). 35 genes are shared between plantiSMASH and MIBiG BGC graphs but absent from any current PlantCyc pathway.

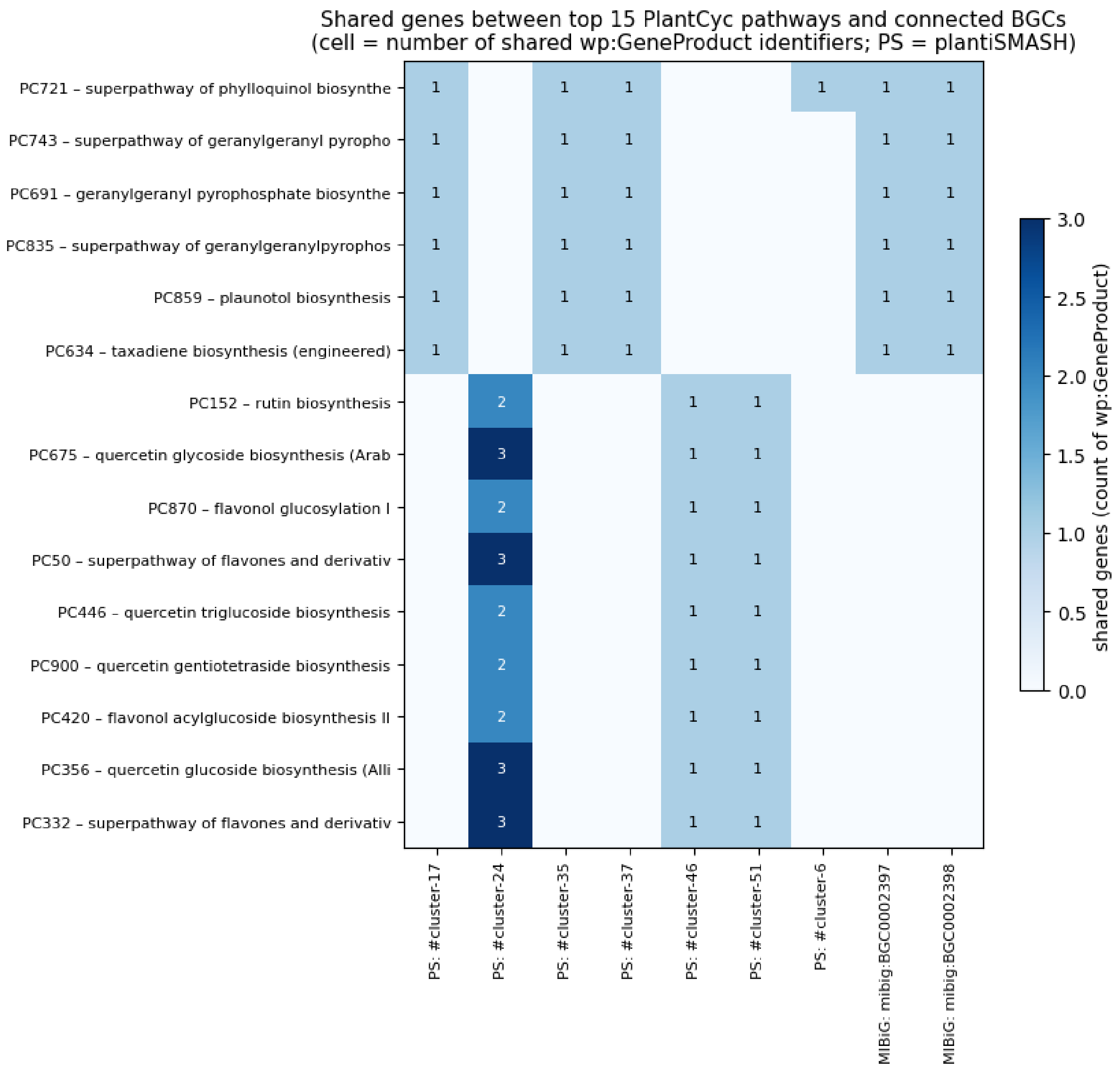


Figure S8. **MIBiG to plantiSMASH BGC coverage representation.** Gene-sharing matrix between the top 15 PlantCyc pathways (ranked by number of connected BGCs) and individual Biosynthetic Gene Clusters from plantiSMASH (PS:) and MIBiG. Rows are PlantCyc pathways labelled by their stable accession (PWY/RXN ID from graph/gpml-properties-extra) and title (truncated to 38 characters). Columns are individual BGC labels. Cell values show the count of shared wp:GeneProduct IRIs (identifiers.org/tair.locus) between a pathway and a BGC; empty cells indicate no shared gene. Full crosslink data (all 225 BGC–gene–pathway triples) in Supplementary Table S3 (bgc_pathway_links_full.csv).

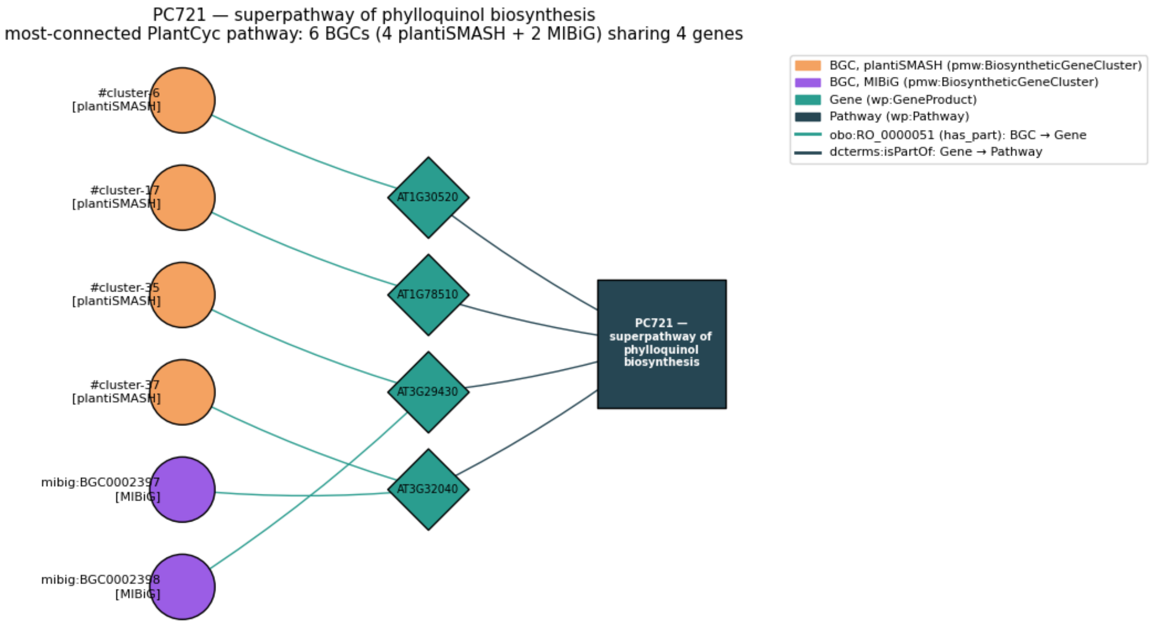


Figure S9. **RDF network of the most-connected PlantCyc pathway** (PWY-5863, *superpathway of phylloquinol biosynthesis*), showing all Biosynthetic Gene Clusters linked to it via shared wp:GeneProduct gene members. Circles (left) represent BGCs (pmw:BiosyntheticGeneCluster), coloured by source: plantiSMASH (orange, *n* = 4) and MIBiG (purple, *n* = 2). Diamonds (centre) represent shared genes (wp:GeneProduct, *n* = 4), labelled by their TAIR locus identifier. The square (right) represents the PlantCyc pathway (wp:Pathway). Green edges: obo:RO_0000051 (has_part, BGC to gene edge); dark edges: dcterms:isPartOf (gene to pathway edge). This tripartite layout illustrates the SPARQL join mechanism between graph/bgc-* and graph/pathways via the shared identifiers.org IRI space: a gene appearing in both a BGC graph and the pathway graph provides the cross-graph link without requiring any pre-computed mapping table.

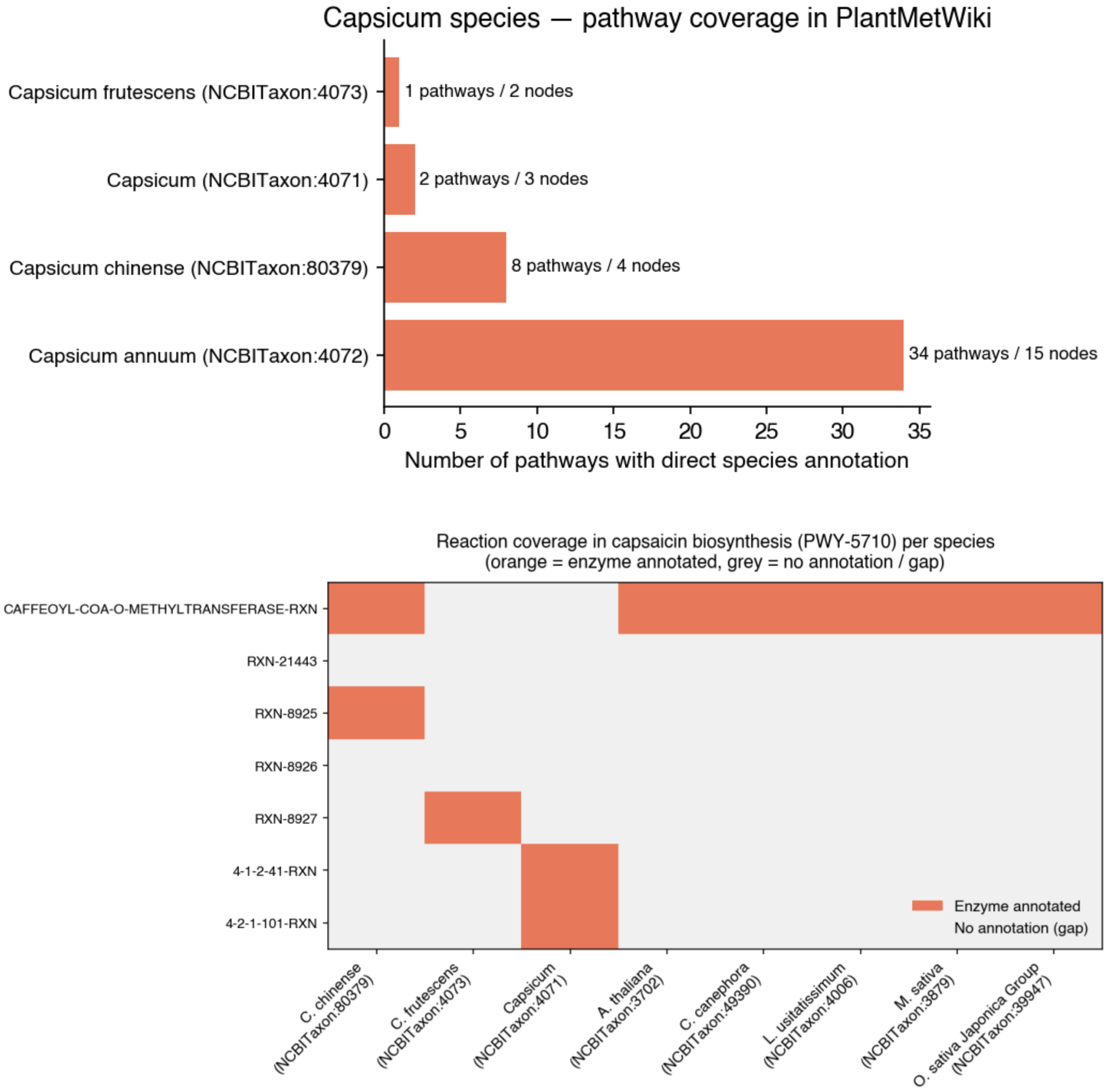


Figure S10. **Top panel: Pathway coverage of *Capsicum* species in PlantMetWiki.** Bars show the number of pathways for which each taxon has at least one directly annotated wp:GeneProduct or wp:Protein DataNode (wp:organism in graph/gpml-taxonomy-extra). The annotation count (number of annotated nodes and pathways) is shown beside each bar. *Capsicum annuum* (NCBITaxon:4072) is the most extensively curated *Capsicum* entry in PlantCyc (CapsicumCyc), covering 34 pathways, while *C. chinense* (NCBITaxon:80379), *C. frutescens* (NCBITaxon:4073), and the genus *Capsicum* (NCBITaxon:4071) have substantially lower annotation depth. **Bottom panel: Reaction coverage matrix for capsaicin biosynthesis** (PWY-5710, *capsaicin biosynthesis*) across all annotated species. Rows are wp:Conversion reactions (PlantCyc reaction accession IDs); columns are species with at least one annotated enzyme in this pathway, labelled with abbreviated binomial name and NCBITaxon identifier. Orange = at least one wp:Protein or wp:GeneProduct DataNode linked via wp:Catalysis; grey = annotation gap. *Capsicum annuum* (NCBITaxon:4072) is absent from this matrix — it has no directly annotated enzyme in this pathway despite being the most pathway-rich *Capsicum* species in PlantMetWiki overall (34 pathways). Cross-species inference candidates are reactions covered in *C. chinense*, *C. frutescens*, or the genus *Capsicum*.

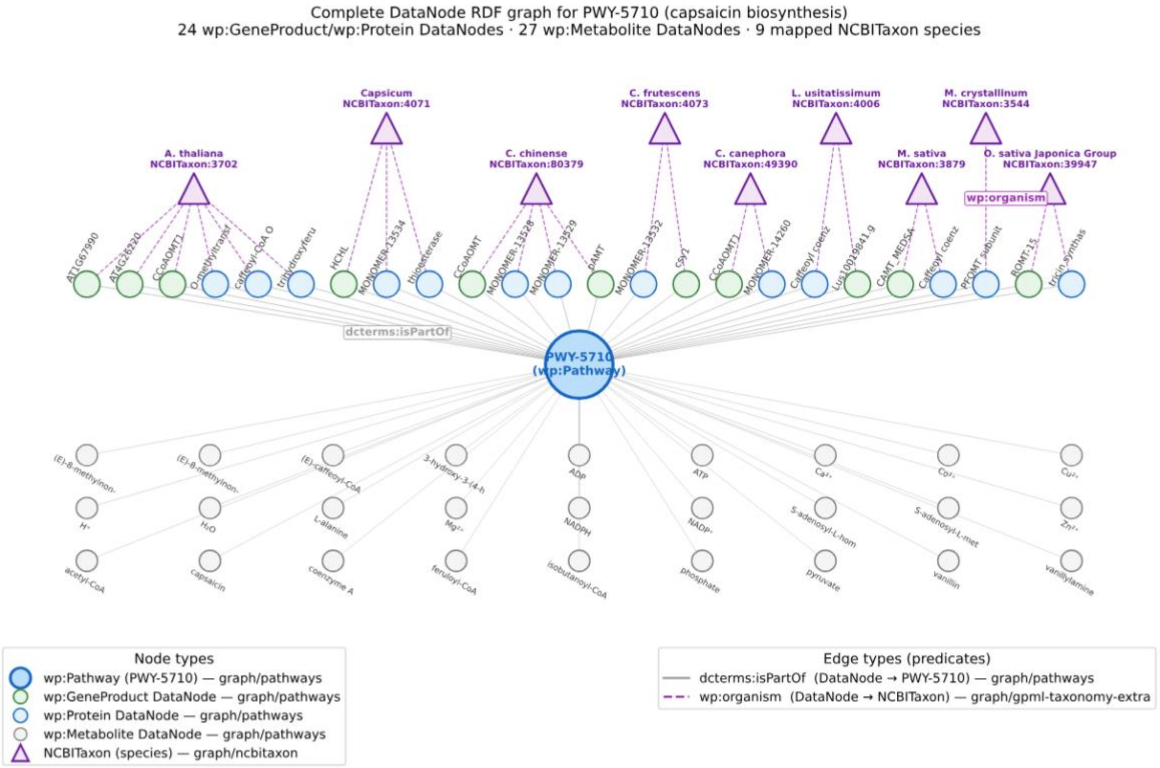


Figure S11. **All DataNodes in capsaicin biosynthesis (PWY-5710) and their species annotations across the two RDF graphs.**
The wp:GeneProduct/wp:Protein DataNodes (green/blue circles) are arranged in a horizontal row, sorted by associated species, and connect to PWY-5710 (centre, wp:Pathway) via dcterms:isPartOf (solid grey edges, graph/pathways). Labels show the DataNode name (up to 14 characters, rotated). The wp:Metabolite DataNodes (grey circles, below) connect the same way: 27 wp:Metabolite nodes, carry no direct species annotation and are shared across all annotated species. NCBITaxon species nodes (purple triangles, above) are labelled with abbreviated binomial name and NCBITaxon accession, connected to their annotated DataNodes via wp:organism (dashed purple edges, graph/gpml-taxonomy-extra). One representative edge label per predicate type is shown for clarity.

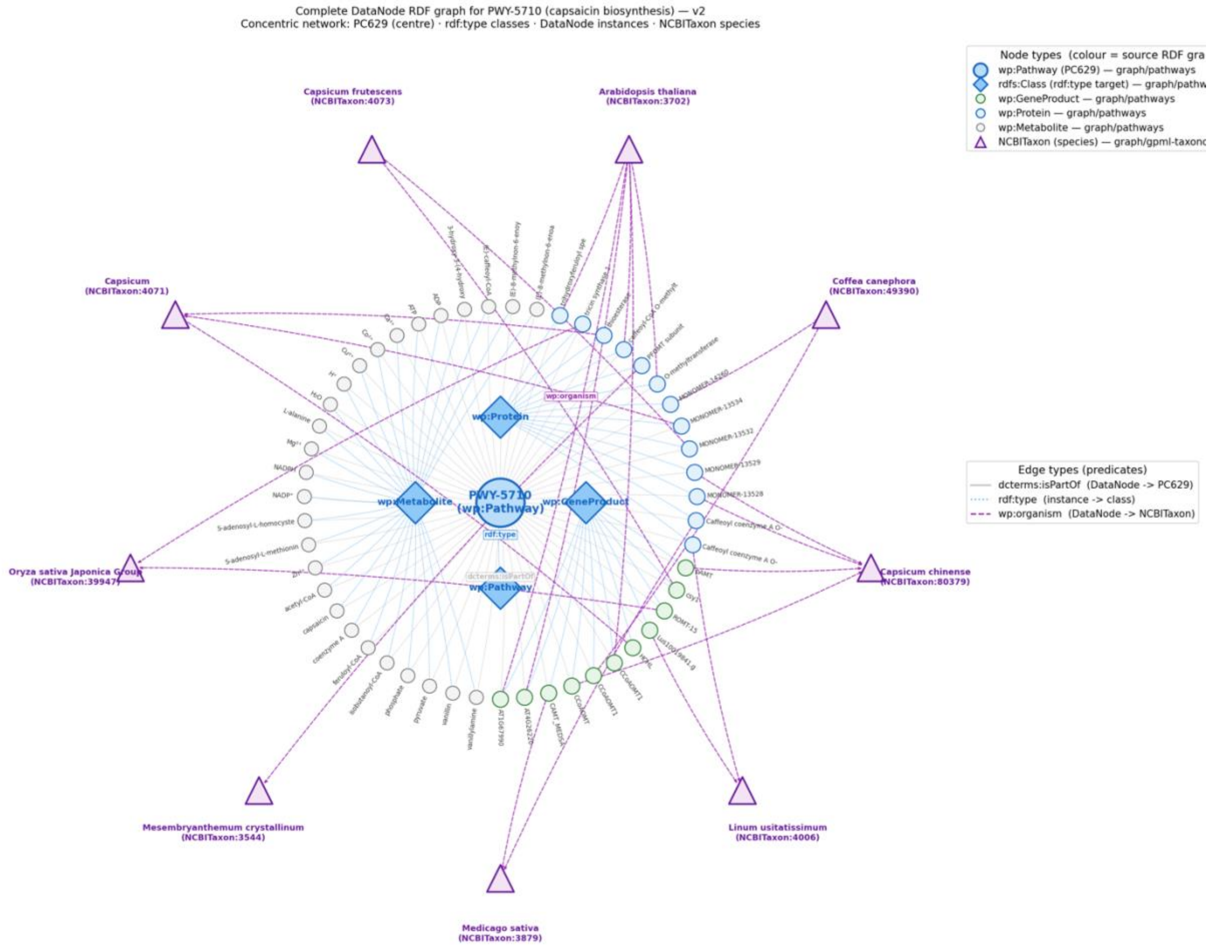


Figure S12. **Complete DataNode RDF graph for capsaicin biosynthesis (PWY-5710) concentric network.** Every node is an individual RDF resource and every edge an individual triple. PWY-5710 (wp:Pathway, large blue circle) sits at the centre; ring 1 holds the four rdfs:Class nodes reachable via rdf:type; ring 2 holds all 51 DataNode instances (11 wp:GeneProduct, 13 wp:Protein, 27 wp:Metabolite), each linked to PWY-5710 via dcterms:isPartOf (grey solid edges) and to its class via rdf:type (blue dotted edges); the outer ring holds the 9 NCBITaxon species nodes, connected to their annotated DataNodes via wp:organism (purple dashed edges, graph/gpml-taxonomy-extra). NCBITaxon nodes are ordered angularly to minimise edge crossings. The data can be exported as Cytoscape.js file for further exploration and editing on Cytoscape Desktop (see Data availability).

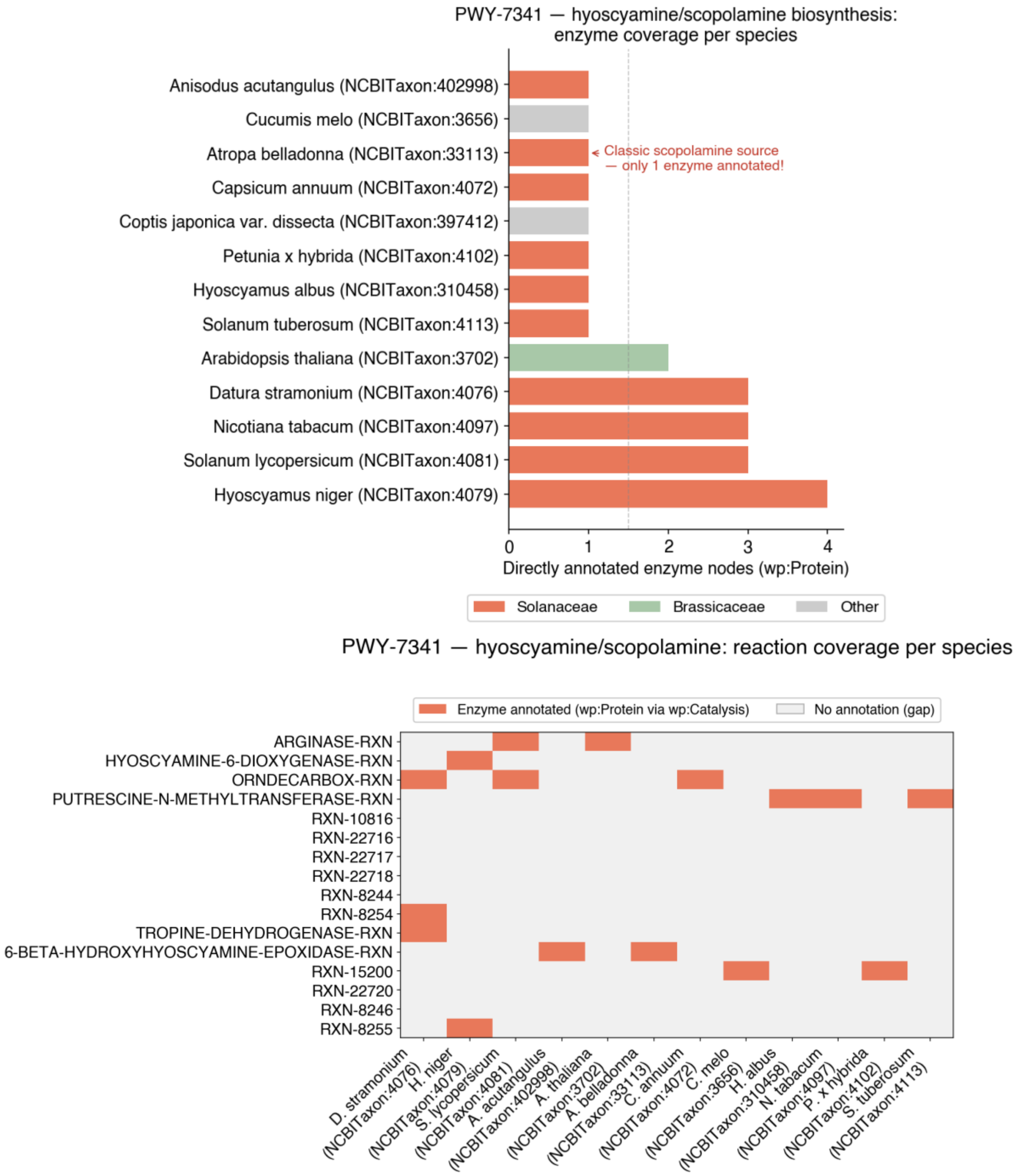


Figure S13. **Top panel: Enzyme coverage per species in hyoscyamine/scopolamine biosynthesis (PWY-7341).** Bars show the count of directly annotated wp:Protein enzyme nodes per species (NCBITaxon ID in parentheses). Orange = Solanaceae species; green = Brassicaceae; grey = other families. *Atropa belladonna* (NCBITaxon:33113), the classical scopolamine source, has only 1 directly annotated enzyme despite 13 species being represented in this pathway, illustrating the curation asymmetry that cross-species inference can address. *Hyoscyamus niger* (NCBITaxon:33116) provides the most complete annotation. **Bottom panel: Reaction coverage matrix for hyoscyamine/scopolamine biosynthesis (PWY-7341)**. Rows are wp:Conversion reactions (PlantCyc accession IDs); columns are species with ≥1 annotated enzyme, labelled with abbreviated name and NCBITaxon ID. Orange = reaction covered by an annotated wp:Protein via wp:Catalysis; grey = gap. Despite *Atropa belladonna* being the principal pharmaceutical source of scopolamine, its annotation depth is far below *Hyoscyamus niger*, making it the primary cross-species inference target in this pathway.

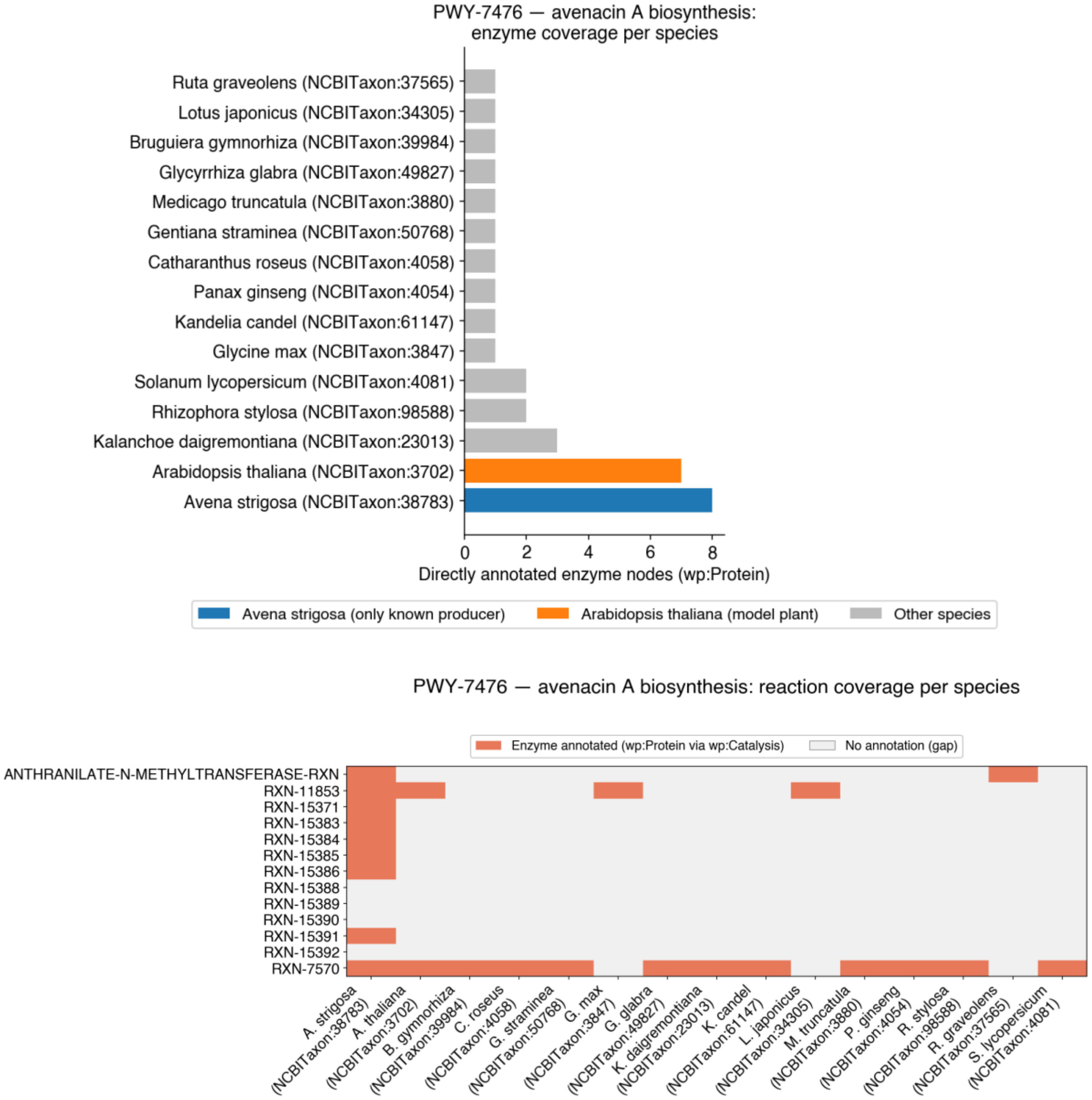


Figure S14. **Top panel: Enzyme coverage per species in avenacin A biosynthesis (PWY-7476)**. Bars show directly annotated wp:Protein enzyme nodes per species (total annotation depth). Blue = *Avena strigosa* (NCBITaxon:38783, the only known producer); orange = *Arabidopsis thaliana* (NCBITaxon:3702, non-producer with orthologous triterpenoid enzymes); grey = other species. Although *Arabidopsis* carries nearly as many annotated enzymes as *Avena* at the enzyme level, at the reaction level it covers only 2 of 13 reactions, both of which are also covered in *Avena* itself.
**Bottom panel: Reaction coverage matrix for avenacin A biosynthesis (PWY-7476)**. Despite *A. thaliana* having comparable enzyme annotation depth to *A. strigosa*, the reaction-level matrix reveals a stark asymmetry: *A. strigosa* covers 9 of 13 reactions while *A. thaliana* covers only 2, both shared with *A. strigosa*. All *A. thaliana* annotation potential is therefore a cross-species transfer candidate, not independent evidence of avenacin production in Arabidopsis.

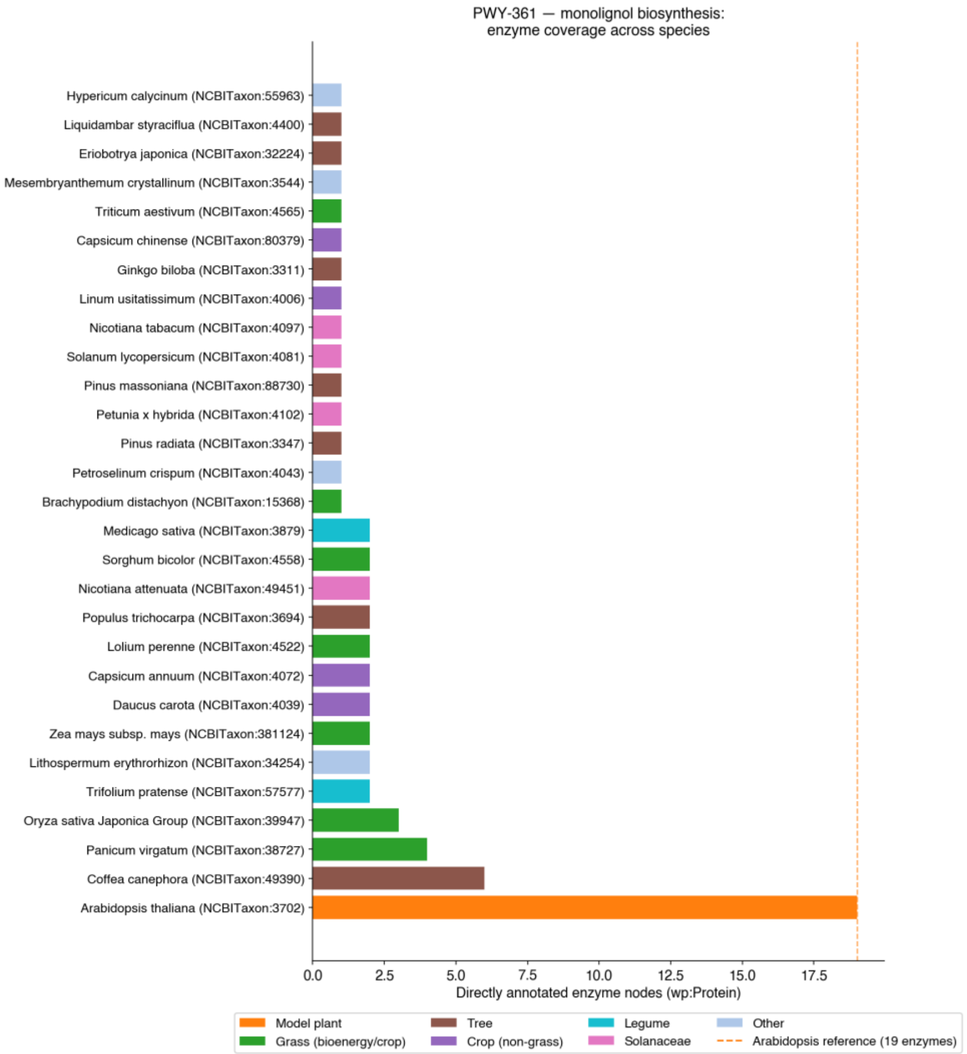


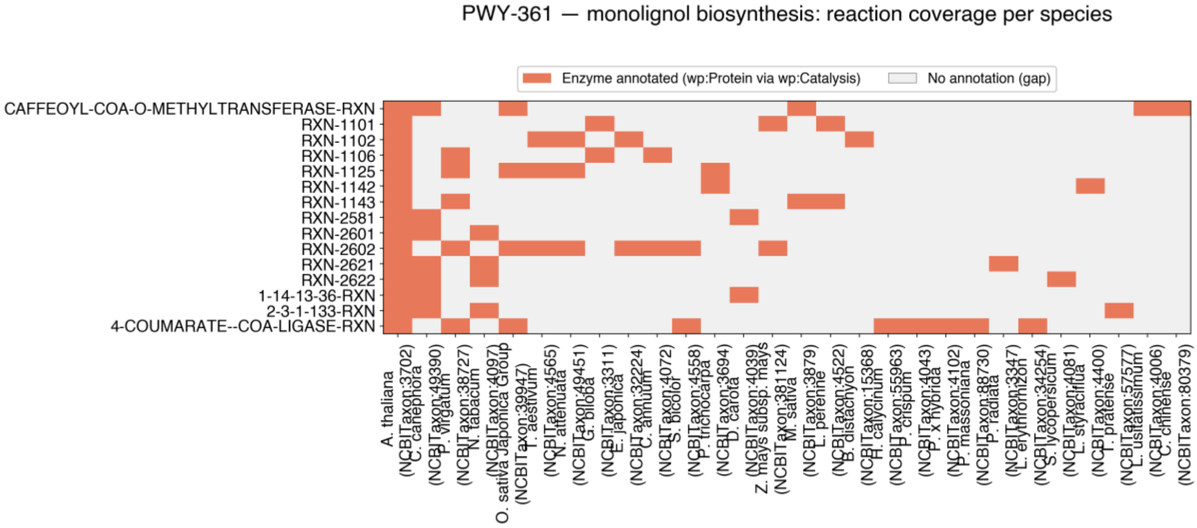


Figure S15. **Top panel: Enzyme coverage per species in monolignol biosynthesis (PWY-361)**, the most taxonomically diverse pathway in PlantMetWiki (29 species). Species are coloured by functional category (model plant, grass, tree, crop, legume, Solanaceae). The dashed vertical line marks *Arabidopsis thaliana*'s (NCBITaxon:3702) reference count of 19 annotated enzymes, the only species covering all 15 reactions in the pathway. Grasses (green bars) show substantially lower coverage, motivating cross-species inference within the Poaceae family.

**Bottom panel: Reaction coverage matrix for monolignol biosynthesis (PWY-361).** *Arabidopsis thaliana* (NCBITaxon:3702) is the only species covering all 15 reactions; all other species show substantial gaps. Within the Poaceae (grasses), the best-covered species reach only 5 of 15 reactions, but combining annotations across the grass sub-family covers 13 of 15, demonstrating the power of cross-family inference for biomass-relevant pathways

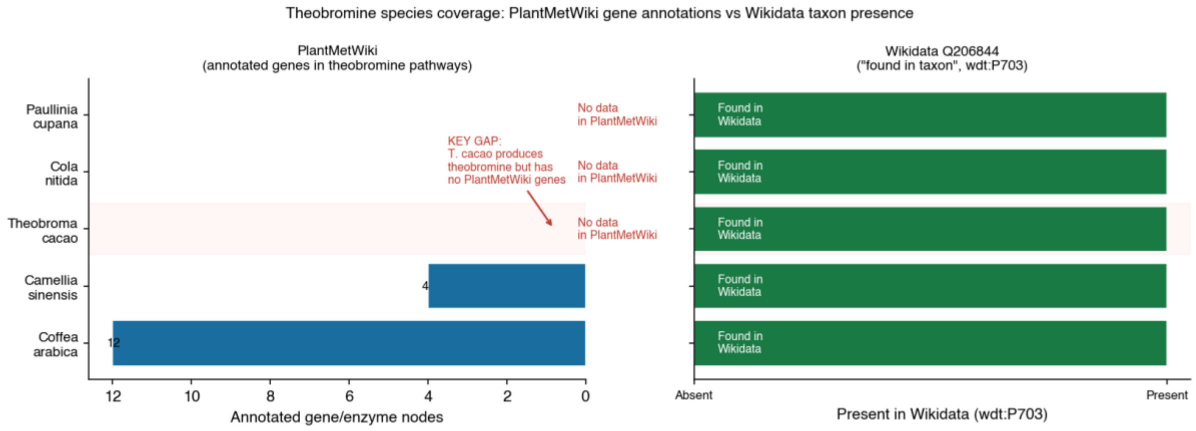


Figure S16. **Cross-species gap analysis for theobromine biosynthesis.** All species known to produce theobromine according to Wikidata (wdt:P703, "found in taxon") shown alongside their PlantMetWiki gene-annotation depth for theobromine-containing pathways. How to read it: species present in Wikidata but with zero/low PlantMetWiki annotation (notably cacao, kola, guarana, coffee and tea are the only well-annotated species) are priority candidates for annotation transfer: Wikidata already provides taxonomic evidence that the pathway should exist in these species.

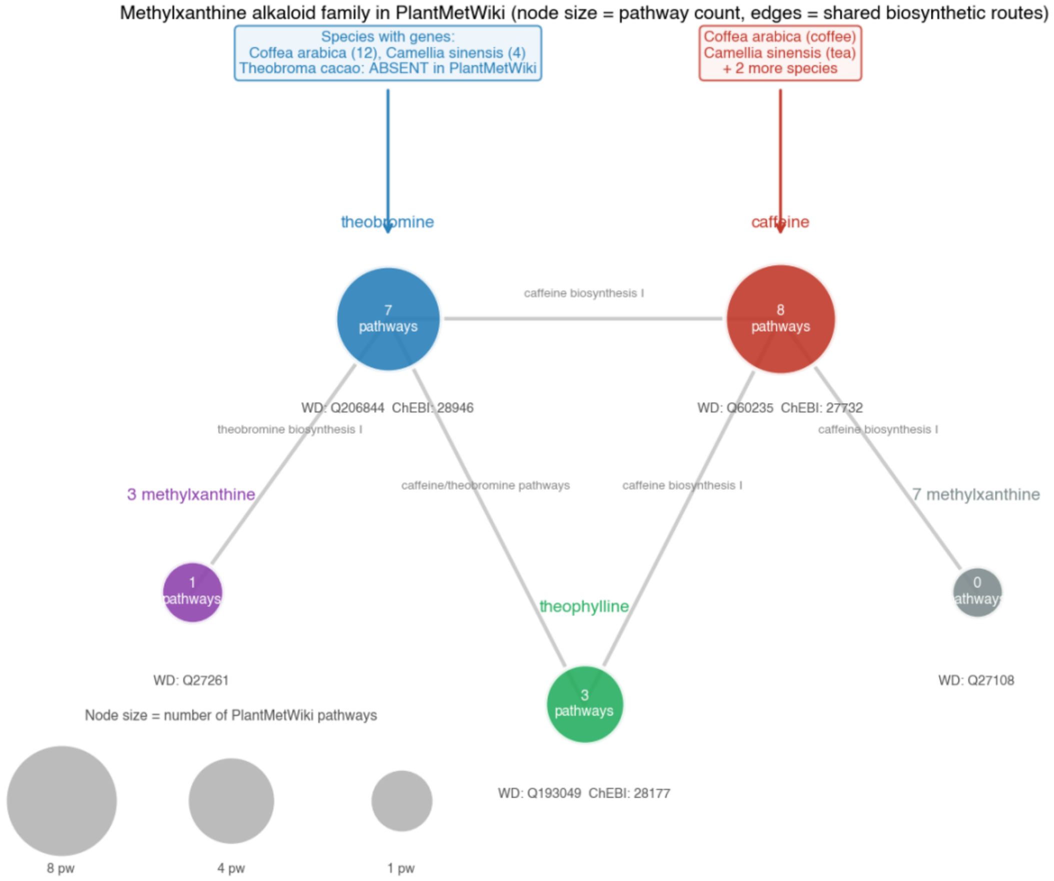


Figure S17. **Methylxanthine metabolic family network.** Caffeine, theobromine, theophylline and related methylxanthines, with node size proportional to the number of PlantMetWiki pathways each compound appears in, and Wikidata QIDs / ChEBI IDs shown as cross-reference labels. How to read it: larger, fully labelled nodes (caffeine: 8 pathways, theobromine: 7) are well-characterised in both PlantMetWiki and Wikidata; smaller nodes (theophylline: 3 pathways, monomethylxanthines: 0) are progressively less curated.

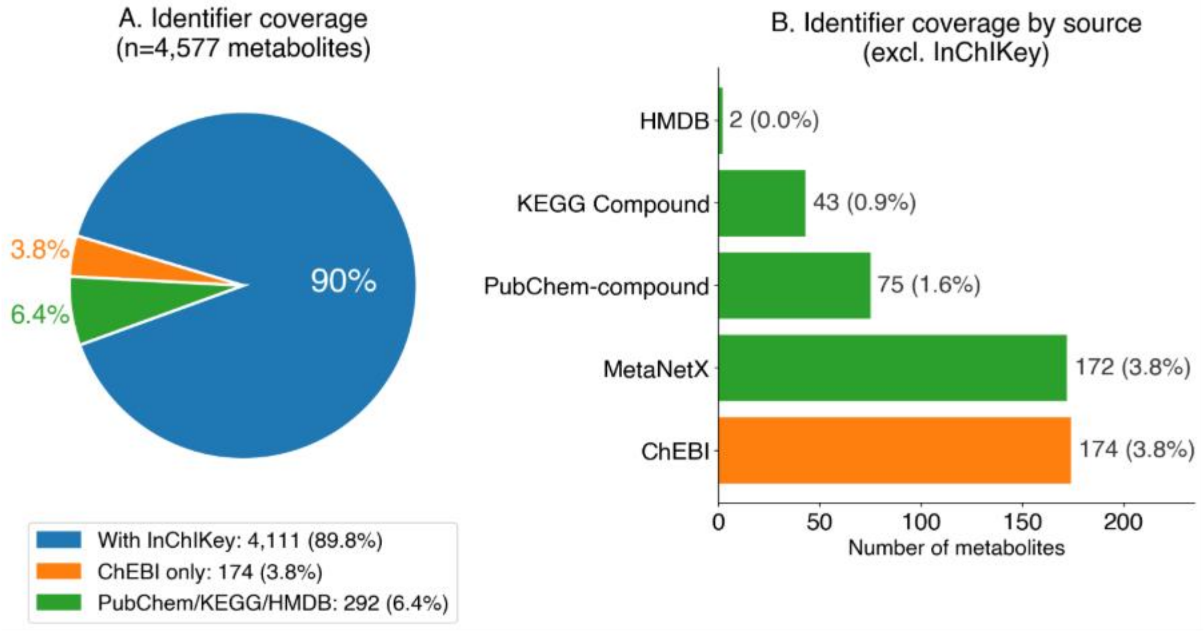


Figure S18. **Metabolite identifier coverage from PlantCyc (before BridgeDB mapping).** (A) Identifier coverage across all 4,577 “wp:Metabolite” nodes in PlantMetWiki: the proportion of metabolites carrying each external identifier type, 90% (4,111) have an InChIKey, the prerequisite for federated SPARQL queries to Wikidata via “wdt:P235”. (B) Breakdown of the other cross-reference databases (ChEBI, MetaNetX, PubChem, KEGG, HMDB) and how many metabolites carry each. As a result, 4,111 metabolites (90%) can be directly federated to Wikidata via InChIKey (wdt:P235).

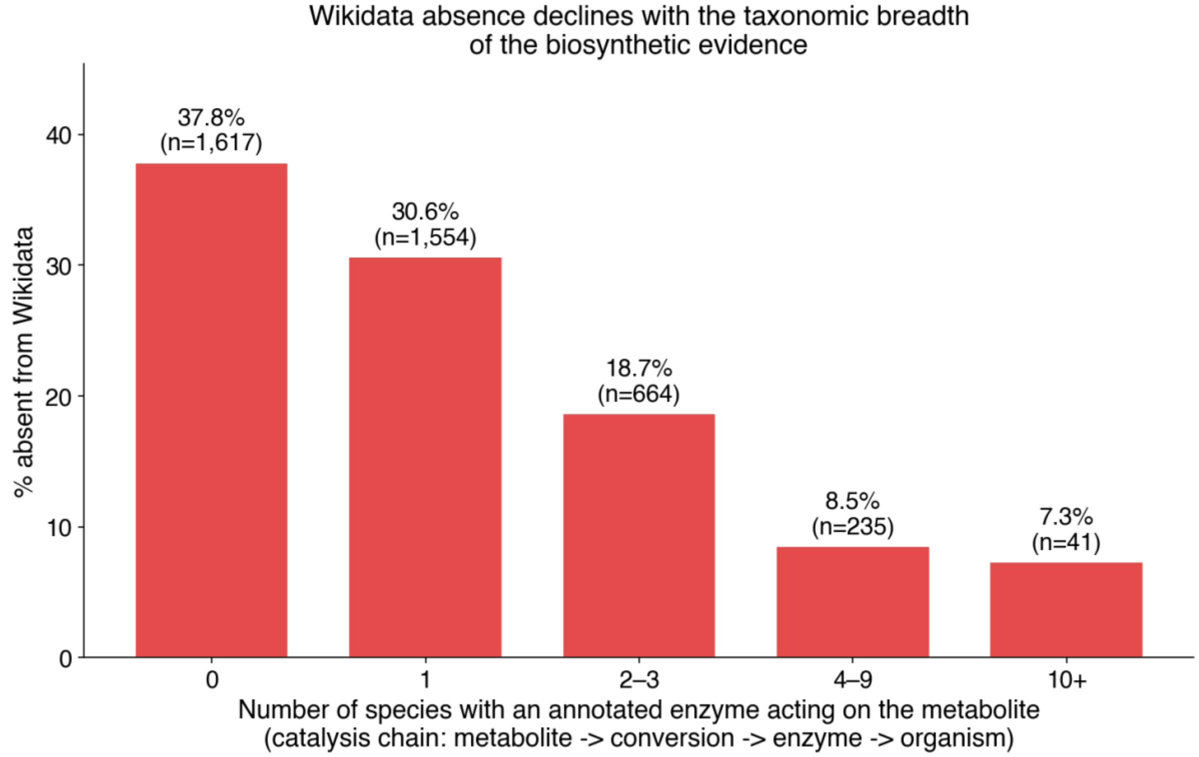


Figure S19. **Wikidata absence declines with the taxonomic breadth of the biosynthetic evidence.** For each of the 4,111 InChIKey-annotated PlantMetWiki metabolites, taxonomic breadth is measured through the catalysis chain: the number of distinct species whose annotated enzyme (wp:Protein, carrying wp:organism) catalyses a wp:Conversion in which the metabolite participates (metabolite → wp:Conversion → catalysing wp:Protein → wp:organism). Bars show the percentage of metabolites in each breadth class that are absent from Wikidata by exact InChIKey; bin sizes (n) are annotated. Absence falls steeply and monotonically, from 37.8% for compounds no annotated species produces (n=1,617), to 30.6% for a single species (n=1,554), 18.7% for two–three (n=664), 8.5% for four–nine (n=235), and 7.3% for ten or more (n=41), showing that the Wikidata gap is concentrated in taxonomically restricted, specialized plant chemistry.
